# Lipoxins Regulate Intercalated Disk-Associated Signaling and Immune Remodeling in Dilated Cardiomyopathy

**DOI:** 10.64898/2026.03.09.710477

**Authors:** Madison Clark, Kyohei Fujita, Lena Anastasia Magdalena Nielsen, Robert T. Johnson, Yusu Gu, Nancy D. Dalton, Bianca Esmée Suur, Ida Bergström, Eric Adler, Ju Chen, Marianne Quiding-Järbrink, Entela Bollano, Niklas Bergh, Matúš Soták, Elisabeth Ehler, Robert Blomgran, Emma Börgeson, Stephan Lange

**Affiliations:** Department of Biomedicine, Aarhus University, Aarhus, Denmark and Steno Diabetes Center Aarhus, Aarhus University Hospital, Aarhus, Denmark; Division of Cardiology, School of Medicine, University of California San Diego, USA; Department of Clinical Immunology and Transfusion Medicine, Sahlgrenska University Hospital, Gothenburg, SE-40530, Sweden; Institute of Advanced Biomedical Engineering and Science, Tokyo Women’s Medical University, Tokyo, Japan; Clinical Department of Clinical Immunology and Transfusion Medicine. Region Östergötland, Linköping, SE-58185, Sweden; Department of Microbiology and Immunology, Institute of Biomedicine, Sahlgrenska Academy, University of Gothenburg, 40530 Gothenburg, Sweden; Department of Cardiology, Sahlgrenska University Hospital, Gothenburg, SE-40530, Sweden; Department of Cardiology, Angereds Närsjukhus, Sweden and Institute of Medicine, Sahlgrenska Academy, University of Gothenburg, Sweden; BHF Centre of Research Excellence at King’s College London, School of Cardiovascular and Metabolic Medicine and Sciences Division & Randall Centre for Cell and Molecular Biophysics, London SE1 1UL, United Kingdom; Department of Biomedical and Clinical Sciences, Linköping University, Linköping, SE-58185, Sweden

**Keywords:** Lipoxin, dilated cardiomyopathy, intercalated disk, muscle lim protein, flow cytometry

## Abstract

We investigated whether pro-resolving lipid mediators of the lipoxin family can attenuate fibrosis and inflammation in muscle LIM protein knockout (MLP^ko^) mice, a model of dilated cardiomyopathy (DCM). Male and female MLP^ko^ mice received either vehicle or a mix of lipoxin-A_4_ and lipoxin-B_4_ three times per week for six weeks. Cardiac function was assessed using echocardiography, and fibrosis and DCM-associated cardiac signaling was evaluated through histology, immunofluorescence and immunoblot analyses. Flow cytometry and RNA sequencing (RNAseq) was performed to identify changes in cardiac gene expression and characterize macrophage subpopulations, respectively. Flow cytometry showed increased inflammatory CD11c^+^ M1-like macrophages and reduction of CD206^+^ M2-like macrophages in MLP^ko^ hearts compared to wild-type controls. Lipoxin treatment partially reversed the macrophage imbalance and showed mild improvements in cardiac physiology in MLP^ko^ males. RNAseq analyses revealed sex-dependent alterations in the expression of pro-fibrotic and inflammation-related genes, suggesting changes in extracellular matrix (ECM) integrity and composition, and to the adaptive immune response. Intriguingly, several ECM proteins showed unexpected localizations at cardiac intercalated disks, which are known to be involved in DCM etiology. Further analysis identified lipoxin-dependent reduction in the DCM-associated expression of intercalated disk components only in lipoxin-treated MLP^ko^ males. Lipoxins also modulated key cardiac signaling pathways in a sex-specific manner, including Erk1/2 and PKCα-linked Ankrd1/Carp1, which is associated with DCM development in MLP^ko^ mice. While lipoxins do not directly reverse cardiac dysfunction or fibrosis in MLP^ko^ mice, they may provide sex-specific protective effects by modulating DCM-related cardiac signaling pathways and by influencing immune-cell populations.

## Introduction

Dilated Cardiomyopathy (DCM) is characterized by progressive ventricular dilation and systolic dysfunction and increased heart failure mortality. The disease affects over 100,000 individuals annually in the US and has an estimated prevalence between 43 to 118 individuals in Western Europe and the US, with a median five-year survival rate despite therapeutical intervention [22, 31, 52, 61]. While DCM has diverse etiologies including aberrant genetic variants in sarcomeric proteins, a unifying pathological feature is ongoing cardiac fibrosis that impairs vascular compliance and contractility [34, 46, 76]. Indeed, cardiac fibrosis represents one of the central drivers of cardiac inflammation and disease progression. Excessive extracellular matrix (ECM) deposition by activated cardiac fibroblasts (myofibroblasts) drives the maladaptive remodeling, yet current heart failure therapies inadequately target the fibrotic process itself [14, 24, 58], necessitating novel therapeutic approaches that directly resolve fibrotic inflammation.

A genetic DCM model that recapitulates many key features of human DCM, including gradual ventricular dilation, systolic dysfunction, and extensive cardiac fibrosis is the muscle LIM protein (MLP/Csrp3) knockout mouse (MLP^ko^) [2]. MLP functions as a sarcomeric mechanosensor required for Z-disc stability, where it interacts with sarcomeric α-actinin, telethonin, and Nrap (nebulin related anchoring protein) to maintain myofibrillar integrity [15, 33, 43, 67]. Loss of MLP triggers both cardiomyocyte stress pathways (fetal gene reactivation, upregulation of the cardiac ankyrin repeat protein Ankrd1/Carp1) and pronounced fibroblast activation, establishing the MLP^ko^ mouse as a robust preclinical model for studying fibrosis-driven DCM pathogenesis [2].

Recent paradigms in inflammation biology emphasize that tissue repair requires active resolution programs mediated by specialized pro-resolving mediators (SPMs) derived from polyunsaturated fatty acids [9, 40]. Lipoxins (LXA_4_ and LXB_4_) are derived from arachidonic acid. Whereas LXA_4_ signals through G protein-coupled receptors (primarily Alx/Fpr2), the receptor for LXB_4_ remains unidentified. Lipoxins limit neutrophil infiltration, promote macrophage efferocytosis, and suppress pro-fibrotic cytokine production [11, 16, 29]. While individual lipoxins have demonstrated anti-fibrotic efficacy in renal and pulmonary models [7, 32], their combined effects in cardiac tissues remain unexplored. Notably, LXA_4_ and LXB_4_ exhibit distinct receptor binding profiles and may provide complementary resolution signals [55], raising the possibility that combined administration may yield complementary pro-resolving effects in the heart. Despite emerging evidence that resolution pathways are dysregulated in heart failure [21], and serum levels of LXA_4_ being upregulated in DCM patients and being proposed as diagnostic disease biomarker [30, 70], previous studies have not examined whether lipoxin-based therapies can attenuate established DCM-associated inflammation and fibrosis, and prevent worsening of the disease. Indeed, evidence from an autoimmune model of myocarditis indicate the therapeutic potential of lipoxins in reducing cardiac inflammation and heart dysfunction [28]. Hence, we hypothesized that combined LXA_4_/LXB_4_ treatment would reduce fibrosis in MLP^ko^ mice through enhanced resolution of cardiac inflammation, thereby improving ventricular function and attenuating molecular markers of heart failure. To test this, we administered lipoxins to MLP^ko^ mice beginning at 4 weeks of age, a time point when fibrosis is established but cardiac decompensation is not yet complete, and assessed physiological, morphological, and molecular outcomes after 6 weeks of treatment.

## Materials and Methods

### Animals and Lipoxin treatment

All procedures were approved by the Animal Care and Use Committee at the University of California San Diego. MLP knockout mice were described previously [2, 37] and maintained on a Black Swiss outbred background (Taconic, Germantown, New York, United States). Both sexes were used; unless otherwise indicated, mice were 4 weeks old at the beginning of the treatment, and 10 weeks at the time of tissue collection. Genotyping of animals was reported elsewhere [37]. Animals were fed standard chow ad libitum and housed at 22°C on a 14 h light/10 h dark cycle. At 4 weeks of age, male and female MLP knockouts were randomly assigned to lipoxin or vehicle treatment groups. Mice received intraperitoneal injections three times per week for 6 weeks with 100 µl of either vehicle (3% ethanol in saline) or a lipoxin cocktail (5 ng/g Lipoxin A₄, #437720-25UG, Millipore, Burlington, Massachusetts, United States; 5 ng/g Lipoxin B₄, #CAYM90420-25, Cayman Chemical, Ann Arbor, Michigan, USA; in 3% ethanol/saline). Lipoxin solutions were prepared on dry ice immediately before injection using molecular biology–grade ethanol (#437433T, VWR) and a 10 µl Hamilton syringe (#20734, Sigma). Body weight was recorded weekly. A schematic of the study design is shown in **Supplemental Figure S1A**.

### Flow cytometry analyses

Antibodies used for flow cytometry analyses are listed in Supplemental Table 1. For antibody titration, freshly isolated cells from spleens were used to determine optimal concentrations. Optimal titers were defined as those giving the brightest positive staining with minimal background and maximal separation between negative and positive populations. Spleens were homogenized through a 70 µm mesh (#352350, Sigma Aldrich) in cold RPMI buffer (RPMI-1640 supplemented with 5% fetal bovine serum and 10 mM HEPES), and cells were resuspended for erythrocyte lysis in 1 ml 1X Lysing Buffer/PharmLyse 10X (#555899, BD Biosciences). Cell counts were obtained to ensure optimal cell numbers for staining, and all subsequent washes were performed in flow cytometry buffer (Ca²⁺/Mg²⁺-free PBS supplemented with 1% heat-inactivated FCS, 0.1% sodium azide, and 1 mM EDTA. 1X Brilliant Stain Buffer (BSD, #566385, BD Biosciences) was added to prevent dye spillover, and cells were blocked with Mouse BD Fc Block (#553141, BD Biosciences) at 4°C. Pre-calculated, titrated antibody volumes were added to each tube, followed by standard wash and incubation steps. Cells were then fixed in 1X Fixative (Stabilizing Fixative 3X Concentrate, #338036, BD Biosciences), washed once more, and incubated overnight at 4°C.

Following titration, flow cytometry analysis of cardiac tissue was performed as described in [54]. Briefly, isolated hearts were immediately perfused via the aorta with 10ml of PBS using a 25-gauge blunted needle. Dissected cardiac ventricles were digested in buffer containing 3 mg/mL Collagenase type II (Worthington, LS004176). Isolated cells were washed in cold PBS + 1% BSA, and cells were resuspended in RPMI to preserve cell integrity. ACK lysis buffer (0.154 M NH₄Cl, 0.009 M KHCO₃, 1.25 M EDTA, pH 7.2) was added to remove erythrocytes. After lysis, cells were washed in flow cytometry buffer and counted to ensure optimal cell numbers for staining.

Cells were transferred to flow cytometry tubes and washed in Ca²⁺/Mg²⁺-free PBS. Live/Dead stain (BD Fixable Viability Stain 510, 564406) was added and incubated for 30 min at room temperature, followed by washes in flow cytometry buffer. Cells were blocked with Mouse BD Fc Block (553141), incubated at room temperature for 5 minutes, then Brilliant Stain Buffer (566385) and a pre-calculated antibody mastermix were added. After a 30 min room-temperature incubation and a final wash in flow cytometry buffer, cells were fixed in 1X Fixative (BD Stabilizing Fixative 3X Concentrate, 338036) and analyzed by flow cytometry using BD ARIA II cell sorter at the Flow Cytometry Research Core Facility located at the VA Hospital in San Diego. Results were analyzed using FlowJo (version 10.10) with a gating strategy shown in Supplemental File 1, and data were plotted as scatterplots to display antibody-positive populations.

### Echocardiography

To assess cardiac function, we employed transthoracic echocardiography as previously described [37]. Briefly, hair on the mouse chest was removed from the left upper abdomen using depilatory cream (Nair). Mice were anesthetized with 1.0–1.5% isoflurane in 0.5 L/min 100% O₂ and maintained under sedation via a nose cone. Animals were placed supine on a heating pad and secured to minimize motion. ECG leads were attached to the upper and lower abdomen for simultaneous electrocardiographic recording. Cardiac function was assessed using a VisualSonics Vevo 2100 with a 55-MHz transducer. M-mode tracings were analyzed for left ventricular internal diameter (LVID) and left ventricular posterior wall thickness (LVPW) at end systole and end diastole. Contractility was expressed as fractional shortening (%FS).

### Transcriptomics

Pre-weighed ventricular tissue was homogenized in QIAzol lysis reagent (79306, Qiagen, Germantown, USA) to generate lysates for RNA purification. Wash and centrifugation steps were performed according to the manufacturer’s instructions to remove contaminants and isolate total RNA. RNA yield was first assessed by NanoDrop, followed by RNA integrity analysis using the Agilent RNA 6000 Nano Kit to confirm suitability for next-generation sequencing of gene expression. For RNAseq analysis, the sequencing library was prepared using a TruSeq stranded mRNA library preparation kit with polyA selection (Illumina). 150 cycles paired-end sequencing were performed using the NovaSeq 6000 system and v1 sequencing chemistry (Illumina) aiming for average coverage of 80M reads per sample. Next generation RNA-Seq was done by the IGM Genomics Center at UC San Diego and the SNP&SEQ-Technology Platform at the University of Uppsala. Bioinformatics analysis was done in Basespace (Illumina) using the RNAExpress tool (using the mouse genome version UCSC mm10) that includes pipelines for sequence alignment (STAR version 2.3.1) and differential expression (DESeq2 version 1.0.17). FDR correction was applied to adjust p values. Comparisons with log2 fold change >0.5 or <-0.5, and -log10(q) > 5 (for wildtype controls vs. MLP^ko^) or -log10(q) > 1.5 (for vehicle vs. LX-treated MLP^ko^) were considered significant.

### Protein analyses

Cardiac ventricles were homogenized, and lysate protein content was normalized based on stain-free gel imaging (Bio-Rad). Equal protein amounts were resolved on 12% SDS-PAGE gels and transferred overnight at room temperature onto nitrocellulose membranes (Biorad). Membranes were blocked in 5% BSA in TBST (10mM Tris-HCl, 0.2% Tween-20, pH 7.4) at room temperature for 2h, then incubated with primary antibodies overnight at 4°C. After three 5 min TBST washes, membranes were incubated with HRP-conjugated secondary antibodies for 1h at room temperature and washed extensively in TBST (10 × 10 min). Proteins were visualized by chemiluminescence using a BioRad ChemiDoc imager. Cardiac actin was used as a loading control for normalization, and quantification was done using ImageJ (NIH, version 1.53) and Excel. A list of primary antibodies used in the study can be found in Supplemental Table 1.

### Immunofluorescence and histology

For immunofluorescence, cardiac tissues were embedded in OCT (Tissue-Tek), frozen, and cryosectioned on a Leica cryostat. Sections were air-dried at room temperature and stored at −80°C until use. Staining was performed as described previously [37]. Briefly, sections were fixed in acetone at −20°C for 5 min, rehydrated in 1× PBS for 5 min, and permeabilized with 0.2% Triton X-100 in 1× PBS for 5 min at room temperature.

Sections were then blocked for 1h in a humid chamber with 5% normal donkey serum (Jackson ImmunoResearch) and 1% BSA (Sigma-Aldrich) in Gold Buffer (55 mM NaCl, 2 mM EGTA, 2 mM MgCl₂, 20 mM Tris-HCl, pH 7.5), followed by overnight incubation at 4°C with primary antibodies diluted in Gold Buffer + 1% BSA. After three 5 min washes in 1× PBS, sections were incubated for 1h with secondary antibodies in Gold Buffer + 1% BSA, washed again three times in 1× PBS, and mounted using Fluorescent Mounting Medium (DAKO). Slides were imaged using a Fluoview 1000 confocal laser scanning microscope (Olympus), equipped with 10x air or 40x oil objective in sequential scanning mode and Zoom rates between 1 and 4. Images were processed using ImageJ (NIH, version 1.53) with the Bio-Formats plugin.

For histology, whole hearts were fixed overnight at 4°C in 4% PFA in 1× PBS. After fixation, hearts were rinsed in 1× PBS for 1 h at room temperature, then incubated overnight at 4°C in 50% OCT diluted in 1× PBS. Tissues were subsequently embedded in OCT, frozen, and cryosectioned on a Leica cryostat. Sections were stained with a hematoxylin and eosin kit (#H-3502-NB, Biotechne) according to the manufacturer’s instructions and imaged using an Olympus SZX12 stereomicroscope equipped with a Lumenera Infinity 3 CCD camera and Infinity Capture software (version 6.4.0, Lumenera).

### Data analysis and availability

Statistical analysis of data was done in Excel (365) and Graphpad Prism (ver. 10.5), with p-values < 0.05 considered as significant. Hierarchical clustering was done in Morpheus (https://software.broadinstitute.org/morpheus), with additional visualization of data using RAWGraphs (https://app.rawgraphs.io/; [50]). Enrichment analysis was done using Metascape (http://metascape.org; [80]). Fully processed next generation sequencing data are shown in Supplemental File 2, while raw data are available upon reasonable request from the authors.

## Results

### Lipoxins reduce DCM-linked cardiac inflammation in male MLP^ko^ mice

Cardiac inflammation is a hallmark of DCM in both patients and animal models [22, 45, 63] . To characterize if MLP^ko^ mice also exhibit signs of cardiac inflammation, and if treatment with lipoxins (LXs; **Supplemental Figure S1a**) can modulate the leukocyte phenotypes associated with the disease, we performed flow cytometry of resident macrophages, neutrophils, B- and T-cells in hearts of wildtype control, vehicle- and LXs-treated MLP^ko^ mice.

Hearts of MLP^ko^ showed a general activation of tissue resident macrophages (CD45^+^LyG6^-^ F4/80^+^CD11b^hi^CCR2^-^Ly-6C^-^CD11a^+^CD11c^+^ cells), with lipoxins having negligible effects compared to vehicle (**Supplemental Figure S1b**). However, male vehicle-treated MLP^ko^ hearts exhibited significant upregulation of inflammatory M1-like macrophages, which was reduced by administration of LXs (p=0.007) (**Figure 1a** left; **Supplemental Table 2**). Surprisingly, female hearts showed reduced M1-like macrophage counts between vehicle or LX-treated mice (p=0.034), compared to wildtype controls, revealing a sex-specific inflammatory response in this DCM model. In contrast, analysis of anti-inflammatory M2-like macrophages revealed significant reduction of cell counts specifically in males (p=0.0001; females: p=0.0514), with LXs showing no effect in the MLP^ko^ hearts regardless of sex (**Figure 1a** center**; Supplemental Table 2**).

**Figure 1.**
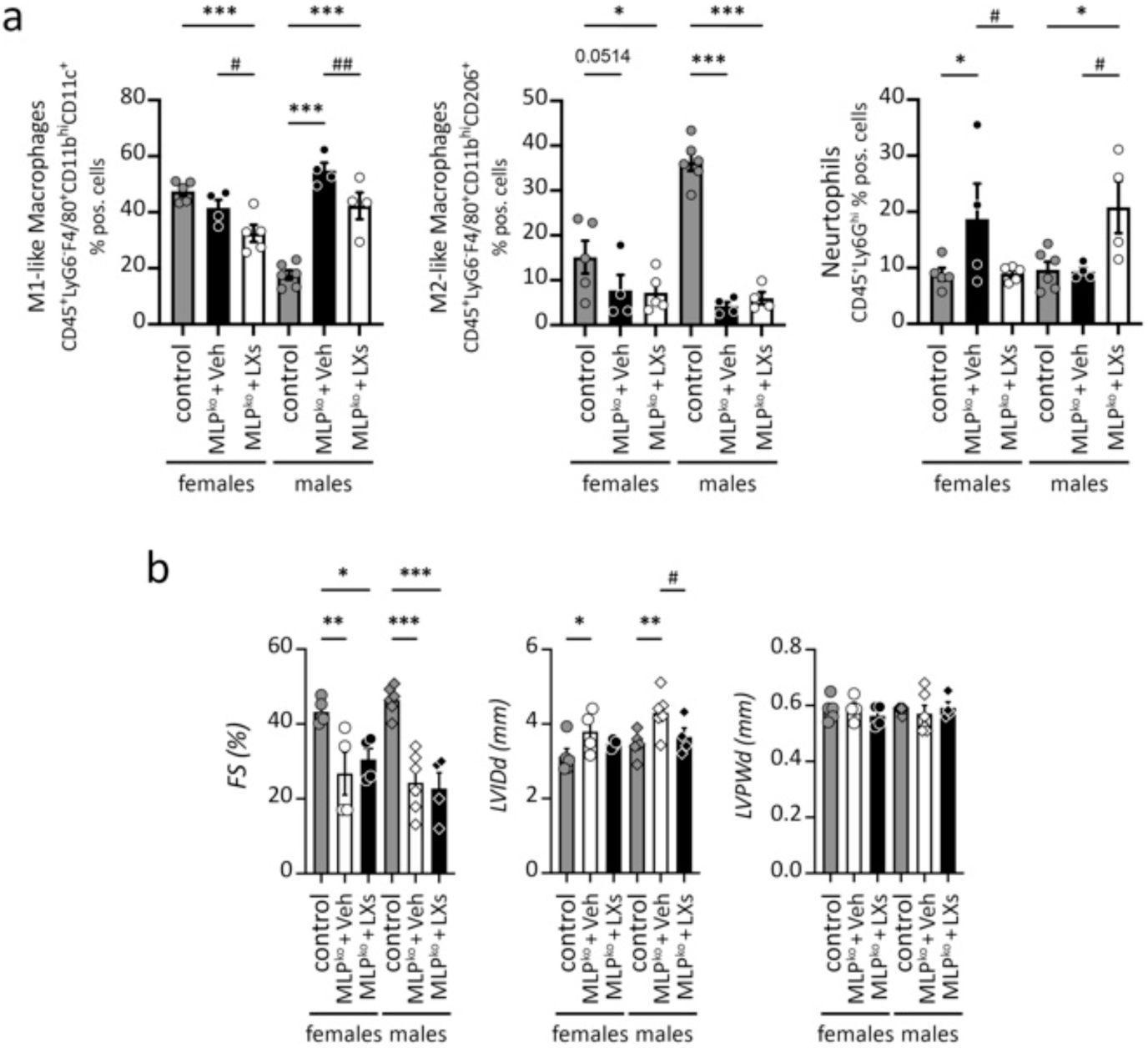
Inflammation and cardiac physiology. **a.** Analysis of cardiac inflammation in 12-week-old wildtype control, vehicle treated MLP^ko^ and LX-treated MLP^ko^ animals. Macrophages were characterized as proinflammatory M1-like macrophages (CD11c^+^) or anti-inflammatory M2-like macrophages (CD206^+^). Data are presented as mean ± SEM. Sample sizes are indicated in the figure, with p-values vs. sex-specific controls (* p<0.05, ** p<0.01, *** p<0.001) or vs. vehicle (# p<0.05, ## p<0.01, ### p<0.001) calculated by ANOVA using Fisher’s LSD test with single pooled variance. **b.** Cardiac physiology determined by transthoracic echocardiography. Shown are fractional shortening (FS, in %; left panel), left ventricular inner dimension during diastole (LVIDd, in mm; middle panel) and left ventricular posterior wall thicknesses during diastole (LVPWd, in mm; right panel) of 12 weeks old male and female control, vehicle treated MLP^ko^ and LX-treated MLP^ko^ mice. Data are presented as mean ± SEM. Sample sizes are indicated in the figure, with p-values vs. sex-specific controls (* p<0.05, ** p<0.01, *** p<0.001) or vs. vehicle (# p<0.05) calculated by ANOVA using Fisher’s LSD test.

Further subclassification indicated that LX treatment of male MLP^ko^ increased counts of recently recruited M1-like macrophages vs. control hearts, while LX treatment of female hearts led to a reduction of recently recruited M2-like anti-inflammatory macrophages compared to vehicle treated MLP^ko^ (**Supplemental Figure S1b**).

Cardiac neutrophil counts were increased in female MLP^ko^, while hearts of male MLP^ko^ showed no changes compared to controls (**Figure 1a** right). Treatment with LXs reduced neutrophil counts in females, but led to a significant increase in male hearts. Looking at strongly activated neutrophils in the recruitment phase, or tissue activated neutrophils, we see low cell counts and no differences between all groups of mice (**Supplemental Table 2**).

We also tested B- and T-cell counts to study changes to the adaptive immune system in DCM. B-cell counts were only reduced in female MLP^ko^ hearts with or without lipoxins compared to controls, while counts decreased in male MLP^ko^ hearts only upon LX treatment (**Supplemental Figure S1c**). Counts of resting/weakly activated tissue resident B-cells declined in MLP^ko^ hearts regardless of sex or LX treatment. Numbers of CCR7^+^, CD11a^+^ or recirculating B-cells (CD45^+^CD45R (B220)^+^CD11a^low^CCR7^+^) remained unchanged between all groups.

Pan T-cell counts were unchanged in all groups, apart from male MLP^ko^ hearts that showed significant increases in T-cell counts, and slight reduction (p=0.08) upon LX treatment (**Supplemental Table 2**). Breakdown of T-cell populations showed increases in CD4^+^ T-helper cells in male and female MLP^ko^ hearts with and without LX, while regulatory T-cells were only elevated in female MLP^ko^ hearts, but returned to control levels upon LX treatment (**Supplemental Figure S1d**; **Supplemental Table 2**). T-killer cell counts were unchanged between all groups of mice, except for significant reduction in LX-treated female MLP^ko^ hearts (vs. controls). CXCR3 positive T-cells were generally low in abundance, except for hearts of control males.

### Treatment with lipoxins reduces left ventricular chamber dilation in male MLP^ko^ mice

Next, we investigated if administration of LXs can improve cardiac functions in MLP^ko^ mice. As expected, transthoracic echocardiography revealed reduced fractional shortening (%FS) and increased left ventricular internal dimension at end diastole (LVIDd) in MLP^ko^ mice compared to wildtype controls, regardless of sex (**Figure 1b**, **Table 1**). Left ventricular posterior wall dimensions in diastole (LVPWd) were unchanged between all groups. Treatment of MLP^ko^ mice with LXs did not improve %FS, compared to vehicle treated controls. However, further analysis of echocardiographic data showed a reduction in LV chamber dilation in LX-treated male MLP^ko^ mice compared to vehicle treated controls. Surprisingly, the effect of LX administration on LV chamber dimensions was not apparent in female MLP^ko^ mice, whose LVIDd remained similar to vehicle treated female controls (**Figure 1b**, **Table 1**).

**Table 1.**
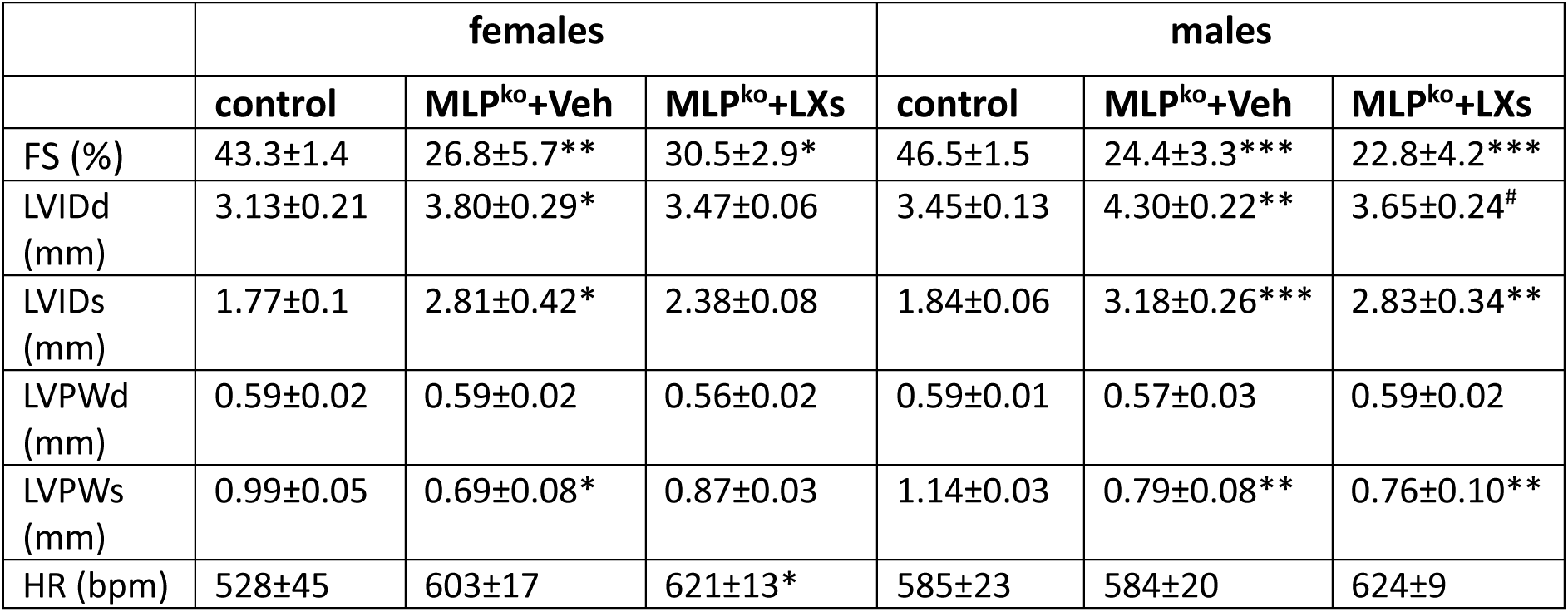
Echocardiography data. Data are presented as mean ± SEM. * p < 0.05, ** p < 0.01, *** p < 0.001 vs. sex-specific controls; # p < 0.05 vs. vehicle treated MLP^ko^, using ANOVA with Fisher’s LSD test.

Additional morphometric analysis of normalized heart (HW/TL) and ventricular (VW/TL) weights (**Supplemental Figure S1e**) showed significant increases in MLP^ko^ females only, when comparing to wildtype control mice. Heart and ventricular weights of male vehicle and LX treated MLPko mice were comparable to those of controls. Atria of vehicle treated male MLP^ko^ mice showed increased weights, albeit with a p-value of 0.061. DCM-associated ventricular chamber dilation could be attributed to altered Hippo-pathway activity [20]. However, when investigated, we only found significant downregulation of YAP in hearts of LX-treated male MLP^ko^ mice, with no further changes to phosphorylation or expression level of additional pathway components, like Sav1, Mob1, Lats1 or Taz (**Supplemental Figures S1f, S1g**). Analysis of lipoxin-treated MLP^ko^ female hearts showed no significant changes compared to their vehicle treated counterparts.

### Transcriptomic profiling reveals sex-specific cardiac remodeling and differential lipoxin responses

To gain deeper insights into molecular mechanisms underlying sex-specific disease phenotypes and therapeutic responses, we performed bulk-RNAseq analysis on ventricle tissue from wildtype, vehicle and LX treated MLP^ko^ mice of both sexes. Pathway enrichment analysis of significantly deregulated genes revealed sex-specific differences between control, vehicle treated and LX-treated MLP^ko^ mouse hearts (**Supplemental Figure S2a**). While muscle specific pathways and mitogen activated protein kinases are enriched in both male and female MLP^ko^ compared to controls, female MLP^ko^ hearts uniquely displayed more terms related to inflammation and stress (e.g. cytokine response, response to heat), compared to male MLP^ko^.

When comparing responses to LX-treatment with vehicle treated MLP^ko^, females showed enrichment in extracellular matrix (ECM)-related pathways (ECM organization, collagen metabolic process, cell migration, supramolecular fiber organization), while the most significant pathway in male MLP^ko^ hearts was related to inflammation (antigen processing, wound healing, IL17A signaling). Comparing overall expression of genes associated with enriched pathways, MLP^ko^ hearts increased overall transcript levels, while treatment with LXs led to a reduction in transcripts compared to vehicle treated MLP^ko^. Across enriched pathways, MLP^ko^ hearts generally showed increased transcript abundance compared to wildtype, whereas LX treatment reduced overall gene expression levels in both sexes, through targeting different pathway sets (**Figure 2a**). Deeper analysis of specific pathways revealed that LX treatment selectively reduced expression of several ECM and fetal gene program markers in female MLP^ko^ hearts (e.g. *Nppa*, *Nppb*; **Figure 2b left panel, Figure 2c**), while males showed no comparable reduction (**Figure 2c right panel**). Hearts of LX treated male MLP^ko^ mice showed changes to expression of genes related to angiogenesis and epithelial cell differentiation and specific reduction of antigen-processing genes (β2-microglobulin [*B2m*], *Psma7* or *Tap1,* **Figure 2b right panel**), the latter of which could also be seen in female hearts (**Figure 2d**). Significantly altered expression of genes involved in antigen processing indicates that infiltration of cardiac tissues by immunoglobulins typically found in DCM [68] is also present in MLP^ko^ hearts. Indeed, study of mouse IgG heavy chain (IgG-HC) levels show increases in expression in MLP^ko^ compared to wildtype controls, specifically affecting female mice (**Figure 2e, Supplemental Figure S2b**). However, treatment of MLP^ko^ with LXs only led to a slight IgG-HC reduction in male hearts (p=0.06), while IgG-HC levels remained elevated in hearts of female MLP^ko^ mice.

**Figure 2.**
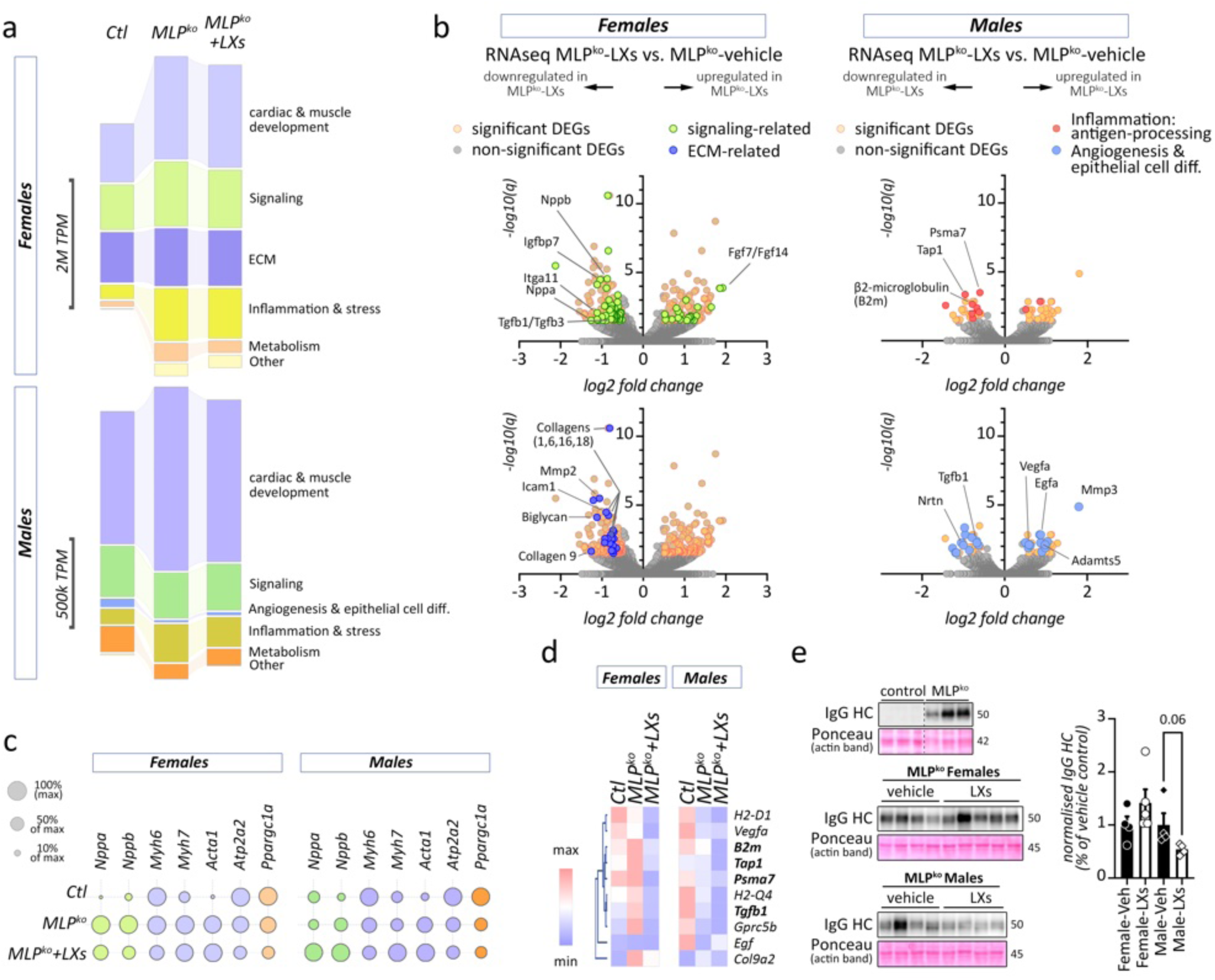
Transcriptome analyses of cardiac hearts and immunoglobulin infiltration. **a.** Analysis of changes to transcriptome levels (in transcripts per million [TPM]) between male and female control (Ctl), MLP^ko^ and lipoxin-treated MLP^ko^ hearts, using significant differentially expressed genes (DEGs) from enrichment analysis categories (Supplemental Figure S2a). **b.** Volcano-plots of extracellular matrix (ECM, blue) and signaling related (green) significant DEGs between vehicle and lipoxin-treated female hearts (left panels), as well as of inflammation (antigen processing, red) and angiogenesis and epithelial cell differentiation-related (blue) significant DEGs between vehicle and lipoxin-treated male hearts (right panels). Other significant DEGs are shown in orange, non-significant DEGs in grey. **c.** Analyses of fetal gene program expression in control (Ctl), MLP^ko^ and LX-treated MLP^ko^ male and female hearts. Shown are relative TPM levels, with natriuretic peptides shown in green, myofilament related genes shown in purple and signaling-related shown in orange. **d.** Heatmap of changes to antigen processing genes in Control (Ctl), MLP^ko^ and LX-treated MLP^ko^ male and female hearts. **e.** Immunoblot analysis of immunoglobulin heavy chain (IgG HC) levels in total cardiac lysates from control (Ctl) and MLP^ko^ hearts (top panel), as well as vehicle and LX-treated MLP^ko^ female (middle panel) and male hearts (bottom panel). Quantification (right) showing mean values ± SEM (in % relative to sex-specific vehicle treated MLP^ko^). Sample sizes are indicated in the figure, with p-values vs. vehicle treated MLP^ko^ calculated by Brown-Forsythe and Welch ANOVA using Welch’s correction.

### DCM-linked interstitial fibrosis is altered by LXs

Given the prominent enrichment of ECM-related pathways in our transcriptome analysis, specifically in the females, we performed comprehensive evaluation of mRNA levels for major ECM structural components and modifying enzymes. Strikingly, female and male MLP^ko^ hearts exhibited diametrically opposed ECM gene expression patterns (**Figure 3a**).

**Figure 3.**
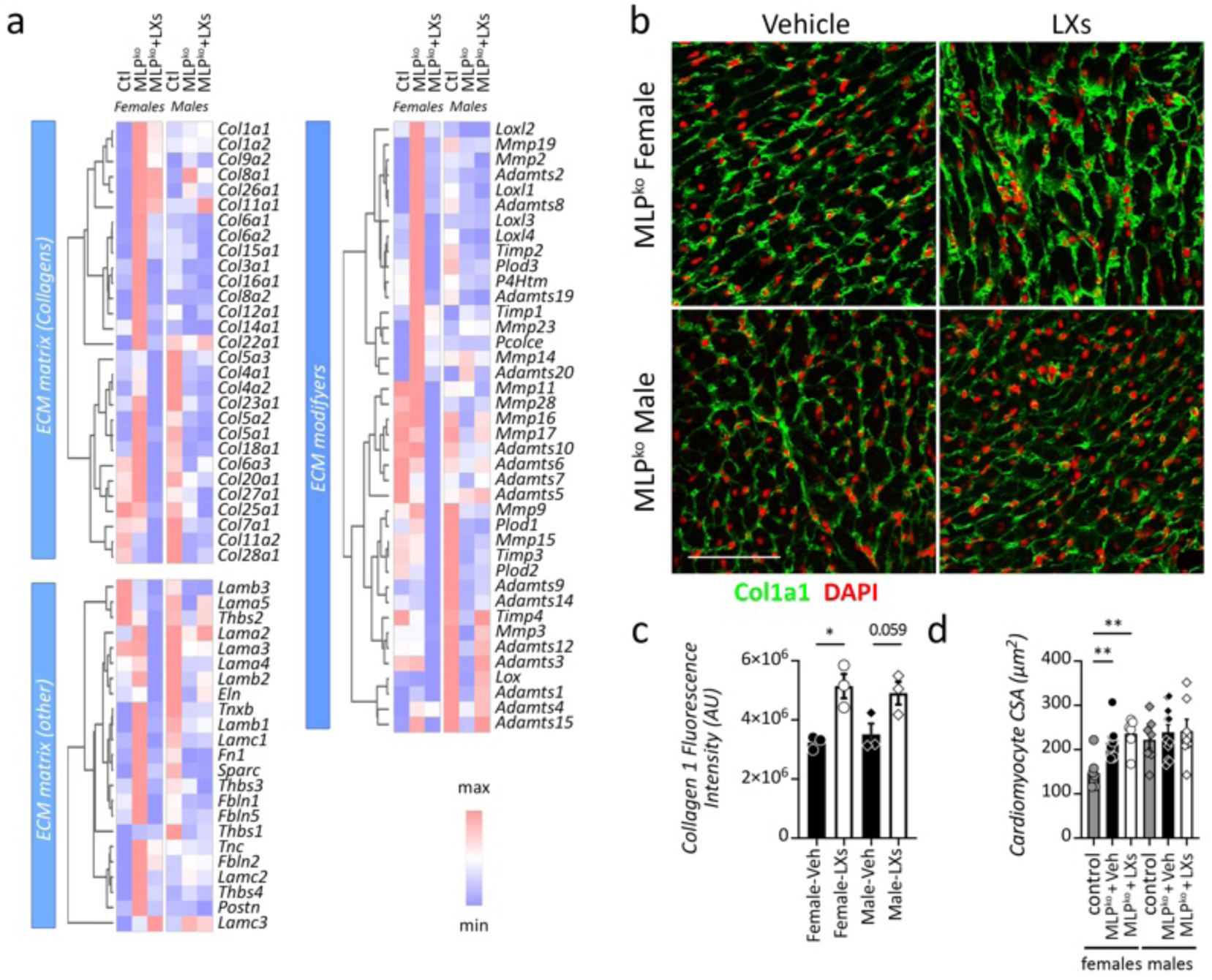
Analysis of cardiac fibrosis and extracellular matrix. **a.** Heatmap of cardiac extracellular matrix (ECM) related mRNA expression levels in Control (Ctl), MLP^ko^ and LX-treated MLP^ko^ male and female hearts. ECM components are subdivided into collagens, ECM modifiers and other ECM matrix components. **b-d.** Immunofluorescence of cardiac cryosections of vehicle and LX-treated MLP^ko^ male and female hearts (b), showing Collagen 1 (Col1a1; green) and DAPI (red). Scalebar = 200µm. Quantification of Col1a1 fluorescence intensity (c) and cardiomyocyte cross-sectional area (d). Data are presented as mean ± SEM, with C showing averages stemming from the analysis of three biological replicates per group. P-values (* p<0.05, ** p<0.01) vs. sex-specific vehicle MLP^ko^ (c) or sex-specific control (Ctl; d) by Brown-Forsythe and Welch ANOVA using Welch’s correction are indicated in the figure.

Female MLP^ko^ hearts displayed widespread upregulation of nearly all matrix genes (e.g. the majority of collagens, *Fn1*, many laminin isoforms and ECM-modifying enzymes, such as *Mmp2*, *Mmp19*, *P4htm*, *Timp1*, *Timp2* or *Pcolce*, compared to hearts of wildtype females (**Figure 3a**). Surprisingly, hearts of male MLP^ko^ mice displayed mostly decreased mRNA levels for almost all ECM matrix components and modifiers, compared to wildtypes, with notable exceptions including matrix components *Col8a1*, *Col26a1*, *Lamc3*, and ECM modifiers *Mmp14*, *Adamts5*, *Adamts14* which were elevated.

Treatment of mice with LXs mostly reverted increased mRNA levels seen in female MLP^ko^ hearts, apart from a cluster of collagen transcripts comprising collagens *Col8a1*, *Col26a1* and *Col11a1*, which remained elevated. In males, LX treatment had minimal effects on mRNA levels of ECM-related genes compared to vehicle treated MLP^ko^ hearts. Transcript levels for several matrix components, including collagens *Col11a1*, *Col22a1* as well as *Eln* and a group of laminins containing *Lama2* and *Lama5* reverted back to levels found in wildtype hearts. LX-treated male hearts also displayed unaltered transcript levels of ECM modifying enzymes, with the exception of *Lox*, *Mmp3*, *Timp4* and several members of the disintegrin and metallopeptidase with thrombospondin type 1 motif (Adamts) gene family, including *Adamts1*, *Adamts3*, *Adamts4*, *Adamts12* and *Adamts15*.

To determine if changes on the mRNA level translate into altered cardiac fibrosis, we studied cardiac sections by histology and immunofluorescence staining. Collagen I protein levels were significantly elevated in MLP^ko^ hearts compared to wildtype controls (**Supplemental Figure S3a-b**). Treatment with LXs resulted in an additional significant elevation of collagen I levels in hearts MLP^ko^ mice compared to vehicle treated controls (p=0.002 vs. p=0.059 in females vs. males, respectively; **Figure 3c**). Masson’s trichrome staining largely corroborated the results from quantification of collagen I (**Supplemental Figure S3c, S3d**). Staining with collagen I also allowed for quantification of cardiomyocyte cross-sectional areas (CSA) across all groups (**Figure 3d**). Loss of MLP only led to significant increases in cardiomyocyte CSA in hearts of vehicle or LX-treated female MLP^ko^, compared to female wildtype controls, with lipoxins showing no effect in preventing CSA increase.

### Enrichment of select ECM components at intercalated disks (ICDs)

Transcript levels of several ECM components, including *Col1a1*, *Mmp9*, *Lox* or *Col11a1*, were specifically altered in hearts of MLP^ko^ mice, and changed again upon LX-treatment. Major collagens like collagen I are typically found on the lateral side of cardiomyocytes, and their localization seemed unaffected between hearts of vehicle and LXs treated MLP^ko^ mice as well as controls (**Figure 3b, Supplemental Figure S3a**). Hence, it was surprising that when evaluating localizations for the ECM modifying enzymes lysyl oxidase (Lox), matrix metallopeptidase 9 (Mmp9) and collagen XI, we noticed either strong or transient localizations at cardiac intercalated disks (ICDs; **Figure 4a-d; Supplemental Figure S4a**). Specifically, Mmp9 and collagen XI showed sex- and treatment-dependent ICD localizations, while Lox displayed strong association to ICDs and unchanged protein levels, regardless of treatment and sex (**Figure 4d, Supplemental Figures S4a, S4b**). Lack of Mmp9 at ICDs in vehicle treated female hearts may be due to reduced levels of the protein in hearts of these mice, compared to the male MLPko mice which exhibited strong Mmp9 ICD enrichment regardless of treatment (**Figure 4b**). Expression of collagen XI was shown to be inversely correlated with Fgf14 levels, with high Fgf14 levels blocking *Col11a1* expression [66]. Intriguingly, *Fgf14* was significantly upregulated only in lipoxin-treated female MLP^ko^ hearts compared to vehicle treated controls (**Figure 2b),** a finding that was corroborated on the protein level (**Figure 4e**). Hence, only male hearts, which show unchanged levels of Fgf14 following LX treatment show collagen XI localization at ICDs, while collagen XI expression is likely blocked by Fgf14 in LX-treated female hearts (**Figure 4c, Supplemental Figure S4a middle panel**).

**Figure 4.**
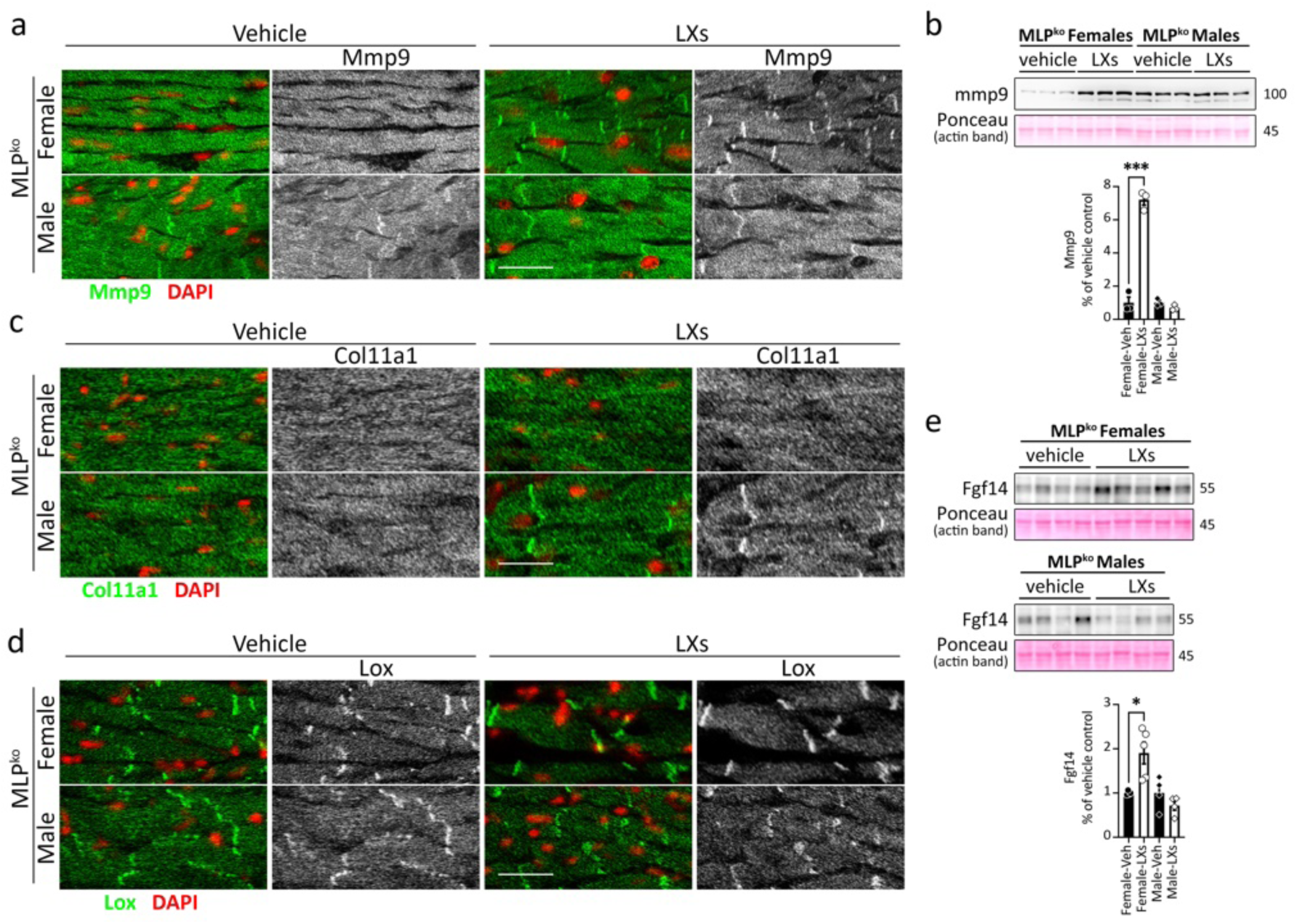
Lipoxin- and sex-dependent intercalated disk localization of ECM components Mmp9, Col11a1 and Lox. **a-b.** Mmp9 localization (green, a) and protein expression levels (b) in hearts of vehicle and LX- treated MLP^ko^ male and female hearts. DAPI (red) is used as counterstain (in a). Ponceau (in b) is shown as loading control. **c-d.** Collagen 11a1 (Col11a1) and Lox localization in hearts of vehicle and LX-treated MLP^ko^ male and female hearts. DAPI (red) is used as counterstain. Scalebar = 20µm (a, c, d). **e.** Fgf14 protein expression levels determined by immunoblot analysis in vehicle and LX-treated MLP^ko^ male and female hearts. Ponceau is shown as loading control. Quantification of protein expression levels (in b and e) is presented as mean ± SEM (in % relative to sex-specific vehicle treated MLP^ko^). Sample sizes and p-values vs. vehicle MLP^ko^ (* p<0.05, ***p>0.001) by Brown-Forsythe and Welch ANOVA using Welch’s correction are indicated in the figure.

Summarized, our finding of ECM components at ICDs is novel and completely unexpected. ICD localization of Lox was independent of sex or treatment. Mmp9 localization at ICDs corresponds in female mice with lipoxin-induced increases in expression of this metallopeptidase, while male MLP^ko^ hearts display ICD localization and expression independent of treatment. However, only male MLP^ko^ mice that have been treated with LXs display collagen XI at the ICD. To our knowledge, β1D integrin is the only ECM associated protein with known ICD localization [5, 69], possibly acting as a receptor and mechanical anchor for collagen XI.

### Improved DCM-linked ICD phenotype in hearts of LX-treated male MLP^ko^

Finding ECM components at ICDs was surprising. However, DCM is known to show characteristic changes at ICDs that alters their structure and signaling, and may contribute to further DCM-development [37]. Indeed, hearts of MLP^ko^ have been characterized to express higher levels of most structural ICD components, like proteins involved in adherens junction formation [15]. We corroborated this finding in immunoblots, which demonstrated significantly elevated levels of all investigated ICD proteins in MLP^ko^ mice when compared to wildtype controls (**Supplemental Figure S5**). Intriguingly, analysis of changes to ICD protein levels revealed that most of the DCM-related upregulation was reverted in hearts of LX-treated male MLP^ko^ mice, when comparing to vehicle treated controls (**Figure 5a, 5b**). Hearts of LX-treated female MLP^ko^ mice showed either unchanged or slight increases over their vehicle treated MLP^ko^ controls.

**Figure 5.**
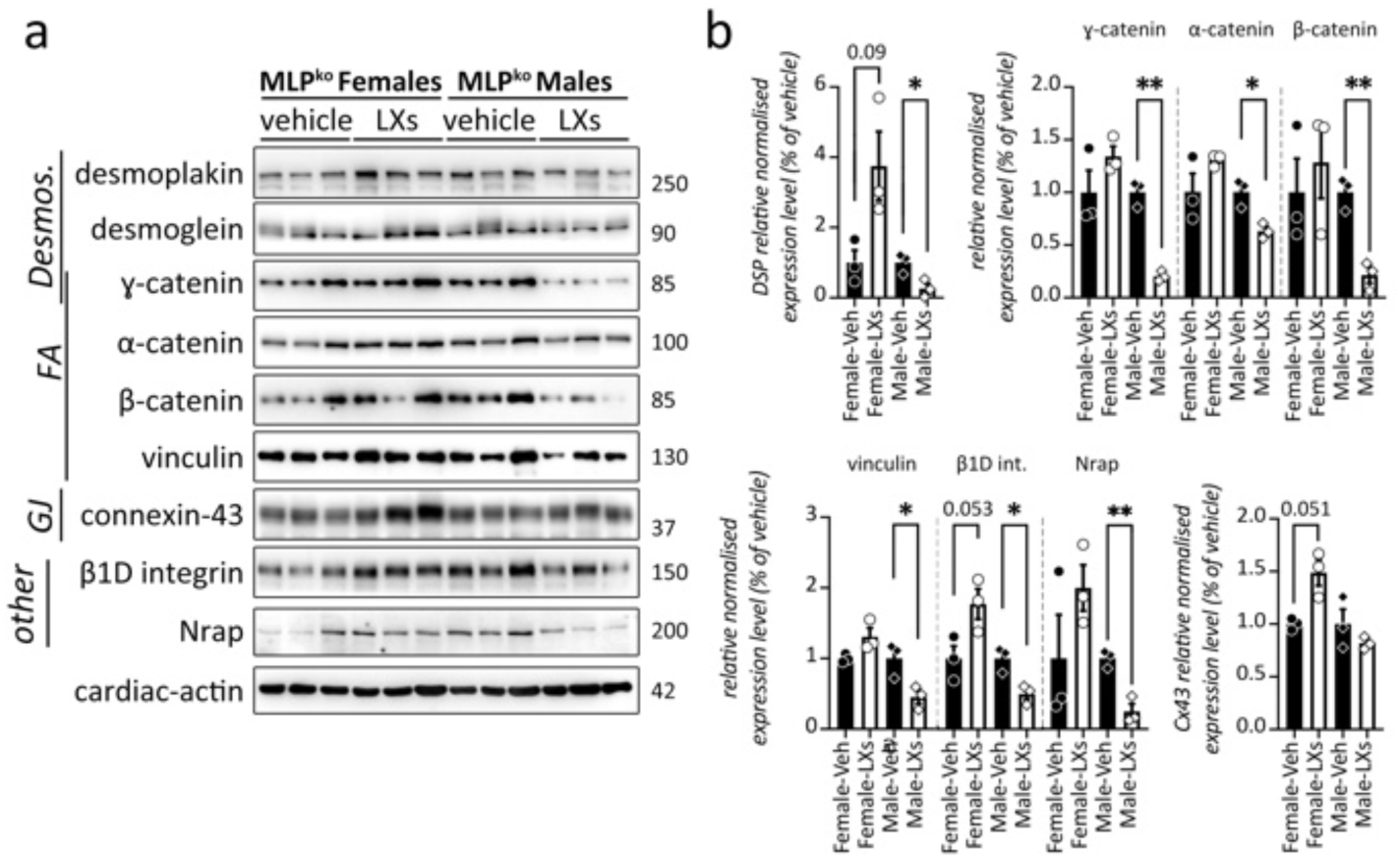
LX reduces pathologically upregulated intercalated disk components in male MLPko mice. **a-b.** Immunoblot analysis (a) and quantification (b) of intercalated disk proteins in vehicle and LX-treated MLP^ko^ male and female hearts. Protein expression levels are presented as mean ± SEM (in % relative to sex-specific vehicle treated MLP^ko^). Sample sizes and p-values vs. vehicle MLP^ko^ (* p<0.05, ** p<0.01) by Brown-Forsythe and Welch ANOVA using Welch’s correction are indicated in the figure.

### LX treatment reduces Ankrd1 levels and pathological kinase signaling in male heart only

Previously, we demonstrated that expression of the cardiac repeat protein CARP/Ankrd1 is linked to pathological PKCα expression and signaling in MLP^ko^ mice, and that genetic ablation of Ankrd1 completely prevents development of DCM in these mice [37]. Given that LX-treatment alleviated chamber dilation and pathological remodeling was alleviated in hearts of LX-treated male MLP^ko^ mice, we wondered whether signaling via Ankrd1 and PKCα may be altered. Indeed, protein levels of Ankrd1 were significantly reduced in LX-treated hearts of male MLP^ko^ only, while females showed unchanged levels of the protein (**Figure 6a, 6b**). Similarly, total PKCα expression was significantly downregulated by LX-treatment in hearts of male MLP^ko^ only (**Figure 6a, 6b; Supplemental Figure S6a**). Beyond total expression, pathological remodeling in DCM was associated with re-localization of Ankrd1 from the sarcomere to ICDs in MLP^ko^, and enrichment of PKCα at ICD structures (**Figure 6c**). Analyzing Ankrd1 and PKCα levels and localization, we noticed increased pathological Ankrd1 enrichment at ICDs in LX-treated female hearts compared to vehicle treated controls (**arrowheads, Figure 6c top panel**), while their male counterparts showed marked redistribution of Ankrd1 back to sarcomeres, where it is normally located [37, 44]. Similarly, PKCα showed increased phosphorylation and stronger association with ICDs in LX-treated female hearts (**arrowheads, Figure 6c bottom panel**), while hearts of male MLP^ko^ mice showed attenuation of the pathological ICD localization of PKCα.

**Figure 6.**
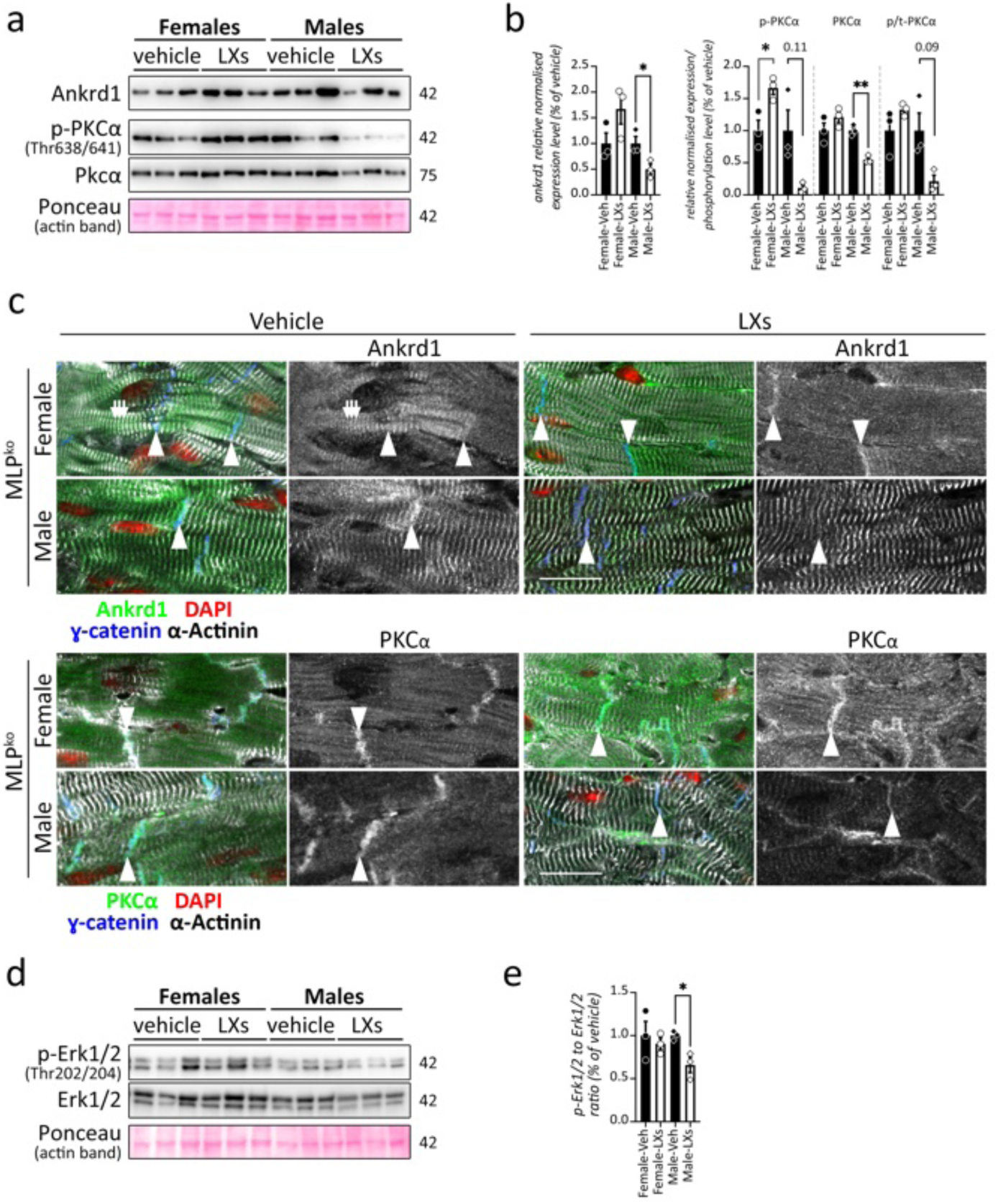
LX alleviates Ankrd1, PKCα and ERK signaling in hearts of male MLPko mice. **a-b.** Immunoblot analysis (a) and quantification (b) of Ankrd1, total and phosphorylated PKCα levels in vehicle and LX-treated MLP^ko^ male and female hearts. Protein expression and phosphorylation levels are presented as mean ± SEM (in % relative to sex-specific vehicle treated MLP^ko^). Sample sizes and p-values vs. vehicle MLP^ko^ (* p<0.05, ** p<0.01) by Brown-Forsythe and Welch ANOVA using Welch’s correction are indicated in the figure. **c.** Immunofluorescence images of frozen cardiac sections stained with antibodies against Ankrd1 (top panel, green in the overlay) or PKCα (bottom panel, green in the overlay). DAPI (red), ɣ-catenin (blue) and sarcomeric α-Actinin-2 (white in the overlay) are used as counterstains. Arrowheads indicate intercalated disks. Scalebar = 20µm. **d-e.** Immunoblot analysis (d) and quantification (e) of total and phosphorylated ERK1/2 levels in vehicle and LX-treated MLP^ko^ male and female hearts. Protein expression and phosphorylation levels are presented as mean ± SEM (in % relative to sex-specific vehicle treated MLP^ko^). Sample sizes and p-values vs. vehicle MLP^ko^ (* p<0.05) by Brown-Forsythe and Welch ANOVA using Welch’s correction are indicated in the figure.

Besides PKC signaling, our RNAseq analysis also indicated changes to MAPK signaling, specifically to the ERK1/2 cascade, as genes associated with this signaling pathway were significantly enriched specifically when comparing changes between wildtype female control and MLP^ko^, as well as vehicle and LX-treated female MLP^ko^ (**Supplemental Figure S2a**). Following these results, we tested expression and phosphorylation levels of Erk1/2, part of the MAPK signaling pathway (**Figures 6d, 6e; Supplemental Figure S6b**). Analysis of phospho/total ratios for Erk1/2 showed significant upregulation in hearts of MLP^ko^ mice (**Supplemental Figure S6b**). Consistent with Ankrd1 and PKCα, phospho/total ratios for Erk1/2 were significantly reduced in hearts of lipoxin treated male MLP^ko^, while their female counterparts showed no improvement (**Figures 6d, 6e**).

Altogether, our data suggest that the reduced ventricular dilation and improved ICD phenotype in lipoxin treated male MLP^ko^ mice, as evidenced by reduced expression of ICD components, is linked to reduced Ankrd1 levels and alleviated activation of PKCα and Erk1/2 kinase signaling. In contrast, LX treatment of female MLP^ko^ mice failed to improve cardiac physiology and pathological ICD remodeling. DCM-associated Ankrd1 expression and localization at ICDs, as well as activation of PKCα and Erk1/2 kinase pathways remained either unchanged or worsened in LX treated female MLP^ko^ hearts. Hence, our findings revealed profound sex specific differences in the therapeutic response to LXs in the heart.

## Discussion

In this study, we set out to investigate if LXs can attenuate DCM in MLP^ko^ mice, a mouse model of the disease. Hearts of LX treated MLP^ko^ mice showed improvements on the physiological and cellular levels by reducing maladaptive extracellular matrix remodeling and cardiac inflammation, alleviating pathological DCM signaling and reducing dilation of the ventricular chamber. However, therapeutic intervention using LXs showed an unexpected sex-specificity: male MLP^ko^ mice showed greater alleviation of DCM symptoms compared to females, with some of the positive outcomes in male mice perhaps being attributable to reduction in pathological signaling via Ankrd1-PKCα signaling pathway [37]. Regardless, improvements of the cardiac physiology were less pronounced compared to disease alleviation achieved by LXA_4_ or its analogue BML-111 in another murine cardiomyopathy model of autoimmune-induced myocarditis [28], also suggesting disease-specific efficacy of lipoxins, and highlighting that only part of the DCM phenotype seen in MLP^ko^ mice may be driven by cardiac inflammation.

### LXs modulate the immune- and fibrotic responses in MLP^ko^

DCM has been demonstrated to increase cardiac inflammation and fibrosis in murine models of the disease and in patients, even in absence of exogenous factors, such as viral mediated myocarditis or bacterial endocarditis [22, 61]. For MLP^ko^ mice it has not previously been demonstrated whether this DCM model also develops increased cardiac inflammation. Our data on significantly increased levels of M1-macrophages in male MLP^ko^, and loss of M2 macrophages in hearts of MLP^ko^ of both sexes validates that this mouse model of DCM recapitulates several aspects of cardiac inflammation seen in patients. Intriguingly, levels of M1 macrophages decreased upon LX-treatment in male mice, suggesting successful targeting of DCM-linked inflammation by systemic administration of LXs. In the cMyBP-C(t/t) mouse model of genetic DCM [45], Lynch et al. also reported a robust increase in M1-macrophages in 3-month-old male mice, whereas M2 macrophage numbers were unchanged, suggesting model-specific changes to cardiac inflammation with a predominantly pro-inflammatory shift at this disease stage. This contrasts with another murine DCM model caused by doxorubicin [59], which resulted in increases of both, M2 and M1 macrophages, the latter being reduced by treatment of mice with melatonin. Notably, both studies used only male mice for their analyses. Our analysis of age-matched MLP^ko^ hearts shows significant M1 activation in males, while females fail to increase in pro-inflammatory M1 macrophages, highlighting a sexually dimorphic inflammatory response in addition to model specific changes, when compared to the cMyBP-C(t/t) and doxorubicin-induced DCM. Moreover, unlike the preserved M2 population in male cMyBP-C(t/t) hearts, we observed a marked loss of resident M2 macrophages in both sexes of MLP^ko^ mice, indicating a broader disruption of macrophage homeostasis in this DCM etiology.

Looking at the innate immune system, we noticed that neutrophil counts only increased in hearts of female MLP^ko^. Typically, neutrophil-leukocyte ratio is used as a predictive marker for DCM [1, 26], with ratios correlating with the heart-failure functional class and pro-NT BNP levels [3]. Surprisingly, male MLP^ko^ mice showed unchanged neutrophil counts when compared to healthy controls. Only LX-treated male MLP^ko^ displayed significantly increases in neutrophils, while LX-treated female MLP^ko^ show a significant reduction. Lower neutrophil counts are typically associated with protection from inflammatory DCM, such as seen in mice with impaired IL17-signalling that is needed for neutrophil recruitment and pro-inflammatory cytokine production [4, 74]. Hence, neutrophil increases in LX-treated MLP^ko^ males, and decreases in LX-treated females are surprisingly not correlating with physiological or molecular changes seen in our analyses, despite enrichment of genes involved in IL17A-signaling in LX-treated MLP^ko^ males (**Supplemental Figure S2a**, bottom right panel). Additional analyses of neutrophil sub-populations did not reveal any further changes.

We also analyzed changes to B- and T-cells of the adaptive immune system, which are known to play important effector and initiator roles for dilated cardiomyopathy development [23, 38]. Total T-cell counts were only elevated in MLP^ko^ males, with LX treatment leading to a slight reduction in their numbers. Looking at CD4^+^ T-helper cells, loss of MLP led to significant increases of cell counts independent of sex, indicating robust adaptive immune-response and T-helper cell activation in DCM. Intriguingly, LXs had no effect on CD4^+^ cell counts, suggesting that underlying mechanisms for their activation are not being targeted by this class of SPMs. CD8^+^ cytotoxic T-killer lymphocyte counts were largely unchanged, with only LX-treated female MLP^ko^ hearts showing a significant reduction compared to healthy controls. Loss of CD8 failed to prevent adverse cardiac remodeling in TAC [38], but played a major role in the initiation of autoimmune-triggered DCM in an animal model of the disease [23]. Thus, unchanged CD8^+^ T-cell counts in LX-treated male MLP^ko^ mice despite improved DCM features, together with the persistence of adverse remodeling in LX-treated female MLP^ko^ mice that show reduction of CD8^+^ cells, suggests that CD4^+^ and CD8^+^ T-cell subsets may not play a central role in the disease progression of this animal model. CD4^+^CD25^+^ regulatory T-cells have been demonstrated to play key roles for autoimmune myocarditis development in a rodent model of the disease [72]. While we noticed a significant increase in cardiac CD45^+^CD3^+^CD4^+^CD127^low^CD25^hi^ regulatory T-cells in MLPko females, generally low levels of this cell-type in the heart, and reduction to levels found in female controls upon LX-treatment suggests negligible function for T-regs in this mouse model of DCM.

Total B-cell counts were lower in female MLP^ko^ regardless of treatment, and LX-treated male MLP^ko^. Analyzing CD11a^+^ adherent/activated B-cells, CCR7^+^ migrating/antigen-presenting B-cells showed no differences in all groups investigated. MLP^ko^ hearts of both sexes however showed a significant drop in resting/weakly activated B-cells, which remained unaltered with LX treatment. Hence, the overall pattern in total B-cells and specific sub-populations suggests a contracting population total of resting B-cell numbers, specifically in LX-treated males, which however maintain high levels of adherent/activated B-cells. B cell-cell interactions have been reported to be increased in DCM, with B-cell-specific CD44 providing extensive interactions with cardiac ECM components [6]. Galectin-9 (Lgals9), which is significantly downregulated in LX-treated female MLP^ko^ hearts was shown to suppress B-cell receptor signaling, suggesting increased B-cell activation specifically in females [19]. This may be reflected by unchanged IgG antibody levels produced by terminally differentiated B-cells in hearts of these mice (**Figure 2e**). LX-treated male MLP^ko^ showed a slight reduction in endomyocardial IgG levels, perhaps indicative of the altered IL17A signaling observed in our RNAseq analysis. However, significant upregulation of recently recruited CD45^+^LyG6^-^F4/80^+^CD11b^hi^CCR7^+^Ly-6C^hi^ macrophages in LX-treated male MLP^ko^ that promote antigen presentation and adaptive immune activation runs counter to this finding.

Cardiac inflammation contributes to activation of resident fibroblasts that drive cardiac fibrosis [17, 25]. While we found that LX treatment decreased mRNA expression levels of many pro-fibrotic and ECM-linked genes (specifically in treated female MLP^ko^ hearts), our histological analyses largely failed to see improvements in interstitial fibrosis in both sexes. In addition, it is unclear what are the causes of the inflammation seen in MLP^ko^ or cMyBP-C(t/t) mouse models of genetic DCM. Given the absence of infections in both models, development of autoantibodies against sarcomeric proteins may play a contributing role in DCM development [22, 79]. Additional research is needed to confirm this.

### LXs affect ICD-localized ECM components and alleviate DCM-linked Ankrd1-PKCα signaling in males

Female MLP^ko^ mice uniquely upregulated *Fgf14* together with reduced expression of fibroblast growth factor receptor 1 (*Fgfr1*). Notably, *Fgf14* were further induced exclusively in female hearts following lipoxin treatment, indicating a sex-specific sensitivity of Fgf signaling to LXs. Unlike canonical FGFs, Fgf14 is intracellular, lacks a signal peptide, is not secreted, does not bind or activate Fgf-receptors, and therefore does not function as a growth factor [73]. Although Fgf14 is thought to be predominantly neuronal, low expression of its two isoforms have been reported in cardiac tissue, albeit only on the RNA level [71]. In neurons, Fgf14 serves as a critical intracellular accessory protein required for Na_v_ and Kcnq2/3 channel expression and function [35, 36, 42, 56, 57, 77]. Aberrant Fgf14 expression is linked to suppression of Col11a1 expression [53, 66]. Consistent with this, our immunofluorescence analyses revealed Col11a1 upregulation exclusively in LX-treated MLP^ko^ males, where Fgf14 is absent, but not in females, which show aberrantly elevated levels of Fgf14.

In our analyses, Col11a1 emerged as one of the few ECM components localized at the ICD, alongside Mmp9 and Lox; a surprising finding given the typical localization of ECM components on the apical side of cardiomyocytes corresponding to the role in making the fibrillar matrix. Increases in Mmp9 are known to be associated with cardiomyopathy [18, 27, 64, 65] and blockage of Mmp9 functions alleviates cardiac remodeling, fibrosis and inflammation [13]. However, its localization to ICD has not been reported. Similar to Mmp9, Lox expression is deregulated in several cardiomyopathy forms, including DCM [64], and genetic variants in Lox were associated with better cardiac function in non-ischemic DCM [51]. Intriguingly, small molecule inhibition of Lox function in a diabetic cardiomyopathy model resulted in improved cardiac function and fibrosis [41], suggesting a potential therapeutic value also in alleviating DCM. ICD localization of Col11a1, Lox and Mmp9 may align with β1D integrin functions [5, 69]. However, while loss of β1D integrin results in DCM [62], its extracellular binding partners at ICDs remain elusive. Remodeling of the ICD is a known driver of DCM pathogenesis in MLP deficiency and includes upregulation of many structural ICD components [15] and the aberrant accumulation and activation of Ankrd1 and PKCα [37]. Lipoxin treatment in male MLP^ko^ mice improved several ICD-associated structural abnormalities, including reducing levels of desmoplakin, α- and γ-catenin, vinculin, and Nrap, accompanied by decreased Ankrd1 and PKCα levels, hallmarks previously linked to disease progression [37]. In contrast, LX-treated female MLP^ko^ hearts displayed the opposite response, with further induction of ICD structural components and increased PKCα phosphorylation, consistent with their exacerbated disease phenotype.

LX-mediated reductions in Ankrd1, PKC, and Erk signaling had few effects on cardiac physiology in either male or female MLP^ko^ mice. We observed significantly reduced left ventricular chamber dilation (LVIDd) only in male MLP^ko^ mice treated with LXs, while fractional shortening remained unchanged in LXs-treated male and females compared with vehicle controls. Unlike MLP × Ankrd1 double-knockout mice, which do not develop DCM [37], the degree of Ankrd1 reduction achieved by LX treatment in male MLP^ko^ hearts is likely insufficient to alter the disease trajectory. Thus, although LXs markedly improved several molecular parameters in MLP^ko^ males and attenuated DCM-associated cardiac inflammation and chamber dilation, LX treatment alone does not suffice to confer meaningful functional rescue, but may contribute to reduce cardiac inflammation in combination with other treatments that counteract the disease, such as β-blockers or Sglt2 inhibitors [22]. Use of NSAIDs and other immunosuppressive or immunoadsorption therapeutic strategies [48, 49, 59] used in the treatment of inflammatory dilated cardiomyopathy (even in absence of viral infection) alongside LXs treatment may also prove beneficial. However, this has not been tested, and represents a limitation of our study.

### Sex-specific effects of LX treatment

An unexpected result of our lipoxin treatment in MLP^ko^ mice were sex-specific effects. Treatment of cell-culture and murine disease models with LXA_4_ and/or LXB_4_ has been done before [7–9, 78], but most experiments were historically done only in male mice. However, few sex-specific effects of LXs have been noted, such as greater Ca^2+^ response in LXA_4_ treated male vs. female conjunctival goblet cells [78]. Differentially regulation of endogenous LXs levels during inflammation were also found to be sex-dependent [39].

Lipoxin A_4_ shares structural features with estrogens (e.g., 17β-estradiol) and can stimulate expression of estrogen responsive genes [10, 47, 60]. LXA_4_ also induced estrogen receptor α phosphorylation, targeting the receptor for degradation by the proteasome [75]. In addition, administration of LXA_4_ increased estrogen receptor β expression in endometriotic stromal cells and suppressed 17β-estradiol-induced p38 MAPK phosphorylation [12]. Other LX precursors, such as arachidonic acid or 15S-HETE as well as LXB_4_ displayed minimal or no binding affinity to the estrogen receptor, confirming the structural specificity [10]. This suggests that sex-specific effects seen in MLP^ko^ mice may be due to LXA_4_ and not LXB_4_ function. However, this represents another limitation of our study. Further experiments are needed to substantiate this theory, and delineate if effects of LX treatment can be attributed to either of the two LXs, or if these represent synergistic effects only seen with the use of both members of this SPM family.

## Conclusions

We demonstrate that lipoxin treatment produces sexually dimorphic molecular and functional effects in genetic dilated cardiomyopathy. In male MLP^ko^ mice, lipoxins reduce cardiac inflammation, normalize ICD protein composition, suppress pathological Ankrd1/PKCα/ERK signaling, and modestly improve ventricular chamber dilation, though without rescuing contractile function. In contrast, female MLP^ko^ mice show no functional benefit and exhibit persistent or exacerbated molecular pathology. These findings reveal unexpected complexity in cardiac resolution physiology and underscore sex as a critical biological variable that must be considered when developing pro-resolution therapies for cardiovascular disease.

## Supporting information

Supplemental File 1 (FACS gating strategy)

Supplemental File 2 (RNAseq data)

## Acknowledgements

The authors would like to thank Andras Harazin, Elia Velasco, Jeppe Høegh Kallesøe and Brigita Medelyte for technical support.

This publication includes data generated at the UC San Diego IGM Genomics Center utilizing an Illumina NovaSeq 6000 that was purchased with funding from a National Institutes of Health SIG grant (#S10 OD026929). The San Diego Flow Cytometry Core is supported by funding from the NIH (P30 AI036214). The authors would like to acknowledge support from Science for Life Laboratory, the National Genomics Infrastructure (NGI), and UPPMAX for providing assistance in massive parallel sequencing and computational infrastructure.

SL is supported by grants from the NIH (HL152251, HL128457), the Swedish Heart Lung Foundation (#20180199), the Novo Nordisk Foundation (NNF22OC0079368), the Aarhus University Research Foundation (AUFF-E-2022-7-9), the Lundbeck Foundation (R396-2022- 189), and the Independent Research Fund Denmark (DFF, #3165-00028B). The work of E.B is support by the European Research Council (ERC-StG no. 804418), Independent Research Fund Denmark (DFF #3165-00026B), Aarhus University Research Foundation (AUFF-E-2022-7-8), Novo Nordisk Foundation (NNF22OC0079363), the Swedish Research Council (VR 2023- 02627), the Swedish state’s ALF-agreement (ALFGBG-978978), and Regionala FoU-medel Västra Götalandsregionen (OLG-2023-02-22). Work in the lab of EE was supported by the British Heart Foundation (RE/18/2/34213) and by UKRI-MRC (MR/R017050/1).

## Statements and Declarations

The authors have no relevant financial or non-financial interests to disclose.

## Author contributions

All authors contributed to the data collection and data analysis. Study design and conception were done by Stephan Lange, Emma Börgeson, Kyohei Fujita and Madison Clark. The first draft of the manuscript was written by Madison Clark, Emma Börgeson and Stephan Lange and all authors commented on previous versions of the manuscript. All authors read and approved the final manuscript.

## Supplemental File 1

Gating strategy and FMO controls for flow cytometry analyses.

## Supplemental File 2

Processed RNAseq data.

---

**Supplemental Table 1.** Antibodies used for flow cytometry and protein analyses.

|  | **antibody/stain** | **clone** | **Manufacturer, catalogue number** |
| --- | --- | --- | --- |
| Flow cytometry | BD Horizon™ Fixable Viability Stain 510 | - | BD, 564406 |
|  | CD11b (Mac-1) | M1/70 | BD, 553312 |
|  | CD206 (Mannose Receptor) | MR5D3 | Thermo Fisher, 53-2061-80 |
|  | F4/80 | T45-2342 | BD, 565410 |
|  | CD45 | 30-F11 | BD, 563891 |
|  | CD192 (CCR2) | 475301 | BD, 747963 |
|  | Ly-6C | AL-21 | BD, 561237 |
|  | CD197 (CCR7) | 4B12 | BD, 740431 |
|  | Ly-6G | 1A8 | BD, 560601 |
|  | CD11a | 2D7 | BD, 742092 |
|  | CD11c | N418 | BD, 565591 |
|  | CD3 | 145-2C11 | BD, 562600 |
|  | CD4 | RM4-5 | BD, 557956 |
|  | CD8 | 53-6.7 | BD, 552877 |
|  | CD45R (B220) | RA3-6B2 | BD, 553087 |
| | CD127 (IL-7 Receptor $\alpha$) | SB/199 | BD, 562419 |
|  | CD25 | PC61 | BD, 557192 |
|  | CD183 (CXCR3) | CXCR3-173 | BD, 562152 |
| Primary antibodies for immunofluorescence and/or immunoblot analyses | Collagen 1 (Col1a1) | E8F4L | Cell Signaling, 72026 |
|  | Mmp9 | D6O3H | Cell Signaling, 13667 |
|  | Mmp9 | - | Proteintech, 10375-2-AP |
|  | Collagen 11a1 (Col11a1) | - | Proteintech, 21841-1-AP |
|  | Collagen 11a1 (Col11a1) | E6A8E | Cell Signaling, 39952 |
|  | Lox | D8F2K | Cell Signaling, 58135 |
|  | Lox |  | Proteintech, 17958-1-AP |
|  | Fgf14 | N56/21 | DSHB, N56/21 |
|  | Desmoplakin | DP2.15 | Sigma, CBL173 |
|  | Desmoglein | M89289 | Biosynth, 10-2298 |
|  | Gamma-catenin/Plakoglobin | - | Cell Signaling, 2309 |
|  | Gamma catenin/Plakoglobin | - | Sigma, SAB2500802 |
|  | Alpha-catenin | - | Proteintech, 13974-1-AP |
|  | Beta-catenin | D10A8 | Cell Signaling, 8480 |
|  | Vinculin | hVIN-1 | Sigma, V9131 |
|  | Connexin-43 | - | Cell Signaling, 3512 |
|  | Beta1D Integrin | - | Developed by Dr. Robert Ross, UC San Diego |
|  | Nrap | - | Developed by Dr. Elisabeth Ehler, King's College London |
|  | Cardiac actin | AC1-20.4.2 | Sigma, A9357 |
|  | Ankrd1 | - | Myomedix, Ankrd1 |
|  | p-PKC-alpha (T638/641) | - | Cell Signaling, 9375 |
|  | PKC-alpha | D7E6E | Cell Signaling, 59754 |
|  | Sarcomeric Alpha actinin-2 | EA-53 | Sigma, A7811 |
|  | p-Erk1/2 (T202/204) | D13.14.4E | Cell Signaling, 4370 |
|  | Erk1/2 | - | Cell Signaling, 9102 |
|  | Sav1 | D6M6X | Cell Signaling, 13301 |
|  | Lats1 | C66B5 | Cell Signaling, 3477 |
|  | Mob1 | E1N9D | Cell Signaling, 13730 |
|  | p-Yap (S127) | D9W2I | Cell Signaling, 13008 |
|  | Yap/Taz | D24E4 | Cell Signaling, 8418 |

**Supplemental Table 2.** Flow cytometry data. Data are presented as mean ± SEM. Statistics (vs. sex-specific control or vehicle when indicated) were done using ANOVA with Fisher’s LSD test.

| Markers |  | Females |  |  | Males |  |  |
| --- | --- | --- | --- | --- | --- | --- | --- |
|  |  | Wildtype Controls | MLP<sup>ko</sup> |  | Wildtype Controls | MLP<sup>ko</sup> |  |
|  |  |  | Vehicle | LXs |  | Vehicle | LXs |
| Mφ<br>(Macrophages) | Pan Macrophages (Mφ)<br>% positive cells | 26.64 ± 1.75 | 36.70 ± 8.75<br>p=0.180 | 29.36 ± 4.41<br>p=0.323 | 48.08 ± 2.45 | 34.30 ± 5.48<br>p=0.061 | 36.85 ± 7.48<br>p=0.742 |
| | Tissue resident Mφ<br>CD45<sup>+</sup>LyG6<sup>-</sup>F4/80<sup>+</sup>CD11b<sup>hi</sup>CCR2<sup>-</sup>Ly-6C<sup>-</sup><br>% pos. cells | 30.46 ± 1.81 | 32.20 ± 9.27<br>p=0.826 | 37.34 ± 7.55<br>p=0.362 | 26.70 ± 1.88 | 28.13 ± 4.87<br>p=0.852 | 33.18 ± 5.56<br>p=0.400 |
| | Activated tissue resident Mφ<br>CD45<sup>+</sup>LyG6<sup>-</sup>F4/80<sup>+</sup>CD11b<sup>hi</sup>CCR2<sup>-</sup>Ly-6C<sup>-</sup><br>CD11a<sup>+</sup>CD11c<sup>+</sup><br>% pos. cells | 77.22 ± 4.82 | 86.20 ± 4.82<br>p=0.094 | **88.36 ± 1.68**<br>**p=0.031** | 56.95 ± 2.61 | **88.33 ± 4.12**<br>**p<0.0001** | **91.98 ± 2.59**<br>**p<0.0001** |
| | Transitional Mφ<br>CD45<sup>+</sup>LyG6<sup>-</sup>F4/80<sup>+</sup>CD11b<sup>hi</sup>CCR2<sup>-</sup>Ly-6C<sup>low</sup><br>% pos. cells | 18.38 ± 2.00 | 10.80 ± 4.02<br>p=0.092 | 11.61 ± 4.52<br>p=0.109 | 7.99 ± 1.52 | 7.77 ± 3.20<br>p=0.958 | 4.92 ± 1.60<br>p=0.467 |
| | Recently recruited Macrophages<br>CD45<sup>+</sup>LyG6<sup>-</sup>F4/80<sup>+</sup>CD11b<sup>hi</sup>CCR7<sup>+</sup>Ly-6C<sup>hi</sup><br>% pos. cells | 5.70 ± 1.25 | 10.32 ± 2.25<br>p=0.349 | 5.31 ± 1.32<br>p=0.932 | 7.45 ± 1.49 | 10.33 ± 1.92<br>p=0.541 | **18.86 ± 8.66**<br>**p=0.022** |
| | CD11c<sup>+</sup> M1-like inflammatory Mφ<br>% positive cells | 47.50 ± 1.65 | 41.55 ± 2.88<br>p=0.152 | **32.48 ± 3.04**<br>**p=0.0006**<br>**(p=0.034 vs vehicle)** | 17.62 ± 1.69 | **54.90 ± 2.73**<br>**p<0.0001** | **42.28 ± 4.76**<br>**p<0.0001**<br>**(p=0.007 vs. vehicle)** |
| | Recently recruited M1-like Mφ<br>CD45<sup>+</sup>LyG6<sup>-</sup>F4/80<sup>+</sup>CD11b<sup>hi</sup>CCR7<sup>+</sup><br>Ly-6C<sup>hi</sup>CD11c<sup>+</sup>CD206<sup>-</sup><br>% pos. cells | 6.89 ± 1.29 | 7.32 ± 2.67<br>p=0.867 | 5.72 ± 1.68<br>p=0.633 | 2.17 ± 0.53 | 6.39 ± 1.68<br>p=0.099 | **11.01 ± 2.91**<br>**p=0.002**<br>**(p=0.101 vs. vehicle)** |
|  | Transitional M1-like Mφ<br>CD45<sup>+</sup>LyG6<sup>-</sup>F4/80<sup>+</sup>CD11b<sup>hi</sup>CCR2<sup>-</sup><br>Ly-6C<sup>low</sup>CD11c<sup>+</sup>CD206<sup>-</sup><br>% pos. cells | 34.84 ± 2.55 | 17.76 ± 4.38<br>p=0.057 | 22.35 ± 10.43<br>p=0.133 | 18.43 ± 2.14 | **36.15 ± 4.10**<br>**p=0.042** | 30.93 ± 7.43<br>p=0.141 |
|  | CD206<sup>+</sup> M2-like anti-inflammatory Mφ<br>% positive cells | 15.14 ± 3.68 | 7.72 ± 3.47<br>p=0.051 | **7.22 ± 1.91**<br>**p=0.029** | 36.35 ± 1.96 | **4.22 ± 0.89**<br>**p<0.0001** | **6.00 ± 1.37**<br>**p<0.0001** |
|  | Recently recruited M2-like Mφ<br>CD45<sup>+</sup>LyG6<sup>-</sup>F4/80<sup>+</sup>CD11b<sup>hi</sup>CCR7<sup>+</sup><br>Ly-6C<sup>hi</sup>CD11c<sup>-</sup>CD206<sup>+</sup><br>% pos. cells | 1.90 ± 1.18 | 5.35 ± 1.67<br>p=0.081 | 0.98 ± 0.81<br>p=0.610<br>**(p=0.031 vs. vehicle)** | 5.37 ± 1.24 | **1.55 ± 0.70**<br>**p=0.046** | 3.84 ± 2.01<br>p=0.408 |
|  | Transitional M2-like Mφ<br>CD45<sup>+</sup>LyG6<sup>-</sup>F4/80<sup>+</sup>CD11b<sup>hi</sup>CCR2<sup>-</sup><br>Ly-6C<sup>low</sup>CD11c<sup>-</sup>CD206<sup>+</sup><br>% pos. cells | 5.33 ± 2.41 | 8.37 ± 6.49<br>p=0.433 | 2.98 ± 1.16<br>p=0.520 | 26.42 ± 1.09 | **4.15 ± 0.74**<br>**p<0.0001** | **4.99 ± 1.27**<br>**p<0.0001** |
| Neutrophils | Pan Neutrophils CD45<sup>+</sup>CD11b<sup>+</sup>LyG6<sup>hi</sup><br>% positive cells | 8.84 ± 1.14 | **18.72 ± 6.31**<br>**p=0.026** | 8.99 ± 0.58<br>p=0.971<br>**(p=0.028 vs. vehicle)** | 9.67 ± 1.42 | 9.34 ± 0.67<br>p=0.936 | **20.83 ± 4.65**<br>**p=0.010**<br>**(p=0.015 vs. vehicle)** |
|  | CD11b<sup>+</sup> Neutrophils<br>CD45<sup>+</sup>LyG6<sup>hi</sup>CD11b<sup>+</sup><br>MFI | 51369 ± 2073 | 45259 ± 3955<br>p=0.204 | 42896 ± 4034<br>p=0.067 | 47879 ± 1745 | 40021 ± 4579<br>p=0.094 | 45965 ± 3183<br>p=0.674 |
|  | Recruitment Phase Neutrophils<br>CD45<sup>+</sup>CD11b<sup>+</sup>LyG6<sup>hi</sup>CD11a<sup>hi</sup>CCR2<sup>+</sup><br>% pos. cells | 15.36 ± 0.34 | 13.63 ± 2.29<br>p=0.795 | 15.39 ± 1.83<br>p=0.997 | 12.75 ± 1.28 | 13.53 ± 2.78<br>p=0.903 | 23.11 ± 12.37<br>p=0.116 |
|  | Tissue activated Neutrophils<br>CD45<sup>+</sup>CD11b<sup>+</sup>LyG6<sup>hi</sup>CD11a<sup>hi</sup>CCR7<sup>+</sup><br>% pos. cells | 4.70 ± 1.02 | 3.25 ± 1.58<br>p=0.516 | 4.35 ± 1.70<br>p=0.868 | 3.54 ± 0.90 | 2.40 ± 1.06<br>p=0.598 | 4.21 ± 2.75<br>p=0.753 |
| B-Cells | B-cells<br>CD45<sup>+</sup>CD45R (B220)<sup>+</sup><br>% pos. cells | 26.54 ± 0.35 | **17.75 ± 3.99**<br>**p=0.039** | **17.15 ± 1.58**<br>**p=0.016** | 27.62 ± 1.72 | 31.03 ± 3.63<br>p=0.386 | **17.87 ± 5.05**<br>**p=0.019**<br>**(p=0.005 vs. vehicle)** |
|  | CD11a<sup>+</sup> adherent/activated B-cells<br>CD45<sup>+</sup>CD45R (B220)<sup>+</sup>CD11a<sup>+</sup> | 88.52 ± 0.77 | 80.25 ± 6.46<br>p=0.344 | 72.58 ± 9.67<br>p=0.051 | 95.95 ± 0.68 | 80.83 ± 4.44<br>p=0.080 | 80.48 ± 3.90<br>p=0.073 |
|  | % pos. cells |  |  |  |  |  |  |
|  | CCR7<sup>+</sup> migrating/antigen-presenting B-cells<br>CD45<sup>+</sup>CD45R (B220)<sup>+</sup>CCR7<sup>+</sup><br>% pos. cells | 5.86 ± 0.61 | 8.76 ± 1.83<br>p=0.103 | 6.46 ± 1.37<br>p=0.698 | 4.85 ± 0.51 | 6.95 ± 1.57<br>p=0.213 | 6.17 ± 0.80<br>p=0.430 |
|  | Resting/weakly activated tissue B-cells<br>CD45<sup>+</sup>CD45R (B220)<sup>+</sup>CD11a<sup>low</sup>CCR7<sup>-</sup><br>% pos. cells | 94.10 ± 0.48 | **89.10 ± 0.93**<br>**p=0.0009** | **89.05 ± 1.15**<br>**p=0.0003** | 94.90 ± 0.50 | **91.38 ± 1.28**<br>**p=0.011** | **91.43 ± 0.75**<br>**p=0.012** |
|  | Recirculating B-cells<br>CD45<sup>+</sup>CD45R (B220)<sup>+</sup>CD11a<sup>low</sup>CCR7<sup>+</sup><br>% pos. cells | 3.57 ± 0.40 | 6.06 ± 1.48<br>p=0.076 | 4.47 ± 1.16<br>p=0.466 | 2.90 ± 0.40 | 4.55 ± 1.09<br>p=0.216 | 3.98 ± 0.49<br>p=0.414 |
| T-cells | Pan T-cells<br>CD45<sup>+</sup>CD3<sup>+</sup><br>% pos. cells | 14.86 ± 1.15 | 18.73 ± 1.57<br>p=0.296 | 17.08 ± 2.21<br>p=0.502 | 9.61 ± 0.68 | **20.33 ± 4.06**<br>**p=0.005** | 13.34 ± 4.60<br>p=0.294<br>(p=0.080 vs.<br>vehicle) |
|  | T-helper cells<br>CD45<sup>+</sup>CD3<sup>+</sup>CD4<sup>+</sup><br>% pos. cells | 21.86 ± 2.75 | **46.55 ± 3.84**<br>**p<0.0001** | **50.63 ± 2.77**<br>**p<0.0001** | 24.28 ± 2.83 | **45.88 ± 4.17**<br>**p<0.0001** | **48.50 ± 2.02**<br>**p<0.0001** |
|  | T-killer cells<br>CD45<sup>+</sup>CD3<sup>+</sup>CD8<sup>+</sup><br>% pos. cells | 45.26 ± 0.81 | 39.00 ± 2.25<br>p=0.116 | **37.37 ± 3.24**<br>**p=0.032** | 42.58 ± 1.79 | 40.90 ± 4.45<br>p=0.653 | 39.00 ± 1.62<br>p=0.342 |
|  | T-regs<br>CD45<sup>+</sup>CD3<sup>+</sup>CD4<sup>+</sup>CD127<sup>low</sup>CD25<sup>hi</sup><br>% pos. cells | 0.55 ± 0.08 | **2.02 ± 0.49**<br>**p=0.0003** | 0.79 ± 0.13<br>p=0.447<br>**(p=0.001 vs.**<br>**vehicle)** | 0.34 ± 0.17 | 0.84 ± 0.21<br>p=0.153 | 0.70 ± 0.34<br>p=0.303 |
|  | CXCR3<sup>+</sup> T-cells<br>CD45<sup>+</sup>CD3<sup>+</sup>CXCR3<sup>+</sup><br>% pos. cells | 1.55 ± 0.28 | 1.56 ± 0.40<br>p=0.998 | 1.10 ± 0.35<br>p=0.884 | 8.25 ± 4.37 | **1.29 ± 0.28**<br>**p=0.043** | **1.19 ± 0.47**<br>**p=0.04** |

**Supplemental Figure S1.**
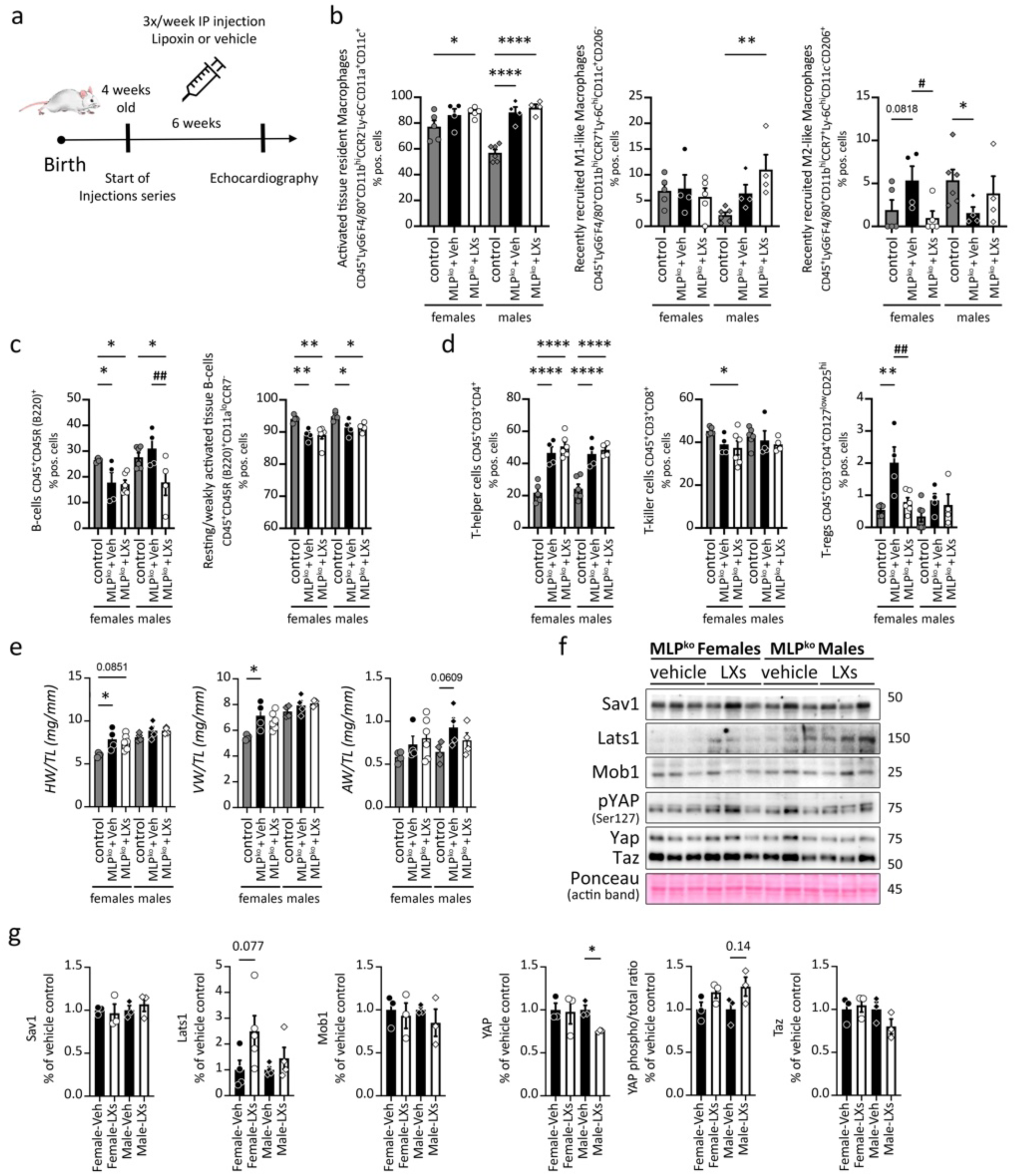
**a.** Study overview. **b-d.** Analysis of macrophage (b), B-cell (c) and regulatory T-cell (d) subpopulations in 12-week-old control, vehicle treated MLP^ko^ and LX-treated MLP^ko^ animals. Data are presented as mean ± SEM. Sample sizes are indicated in the figure, with p-values vs. vs. sex-specific controls (* p<0.05, ** p<0.01, *** p<0.001, **** p<0.0001) or vehicle treated mice (# p<0.05, ## p<0.01) calculated by ANOVA using Fisher’s LSD test with single pooled variance. **e.** Morphometric analyses of heart weight to tibia lengths (HW/TL) ratios, ventricular weight to tibia lengths (VW/TL) ratios and atria weights to tibia lengths (AW/TL) ratios. Data are presented as mean ± SEM. Sample sizes and p-values vs. vs. sex-specific vehicle MLP^ko^ (* p<0.05) by ANOVA using Fisher’s LSD are indicated in the figure. **f-g.** Immunoblot analysis (f) and quantification of Hippo-signaling pathway component protein expression levels (g). Ponceau is shown as loading control. Data are presented as mean ± SEM (relative to sex- specific vehicle treated MLP^ko^). Sample sizes and p-values vs. vehicle MLP^ko^ (* p<0.05) by Brown-Forsythe and Welch ANOVA using Welch’s correction are indicated in the figure.

**Supplemental Figure S2.**
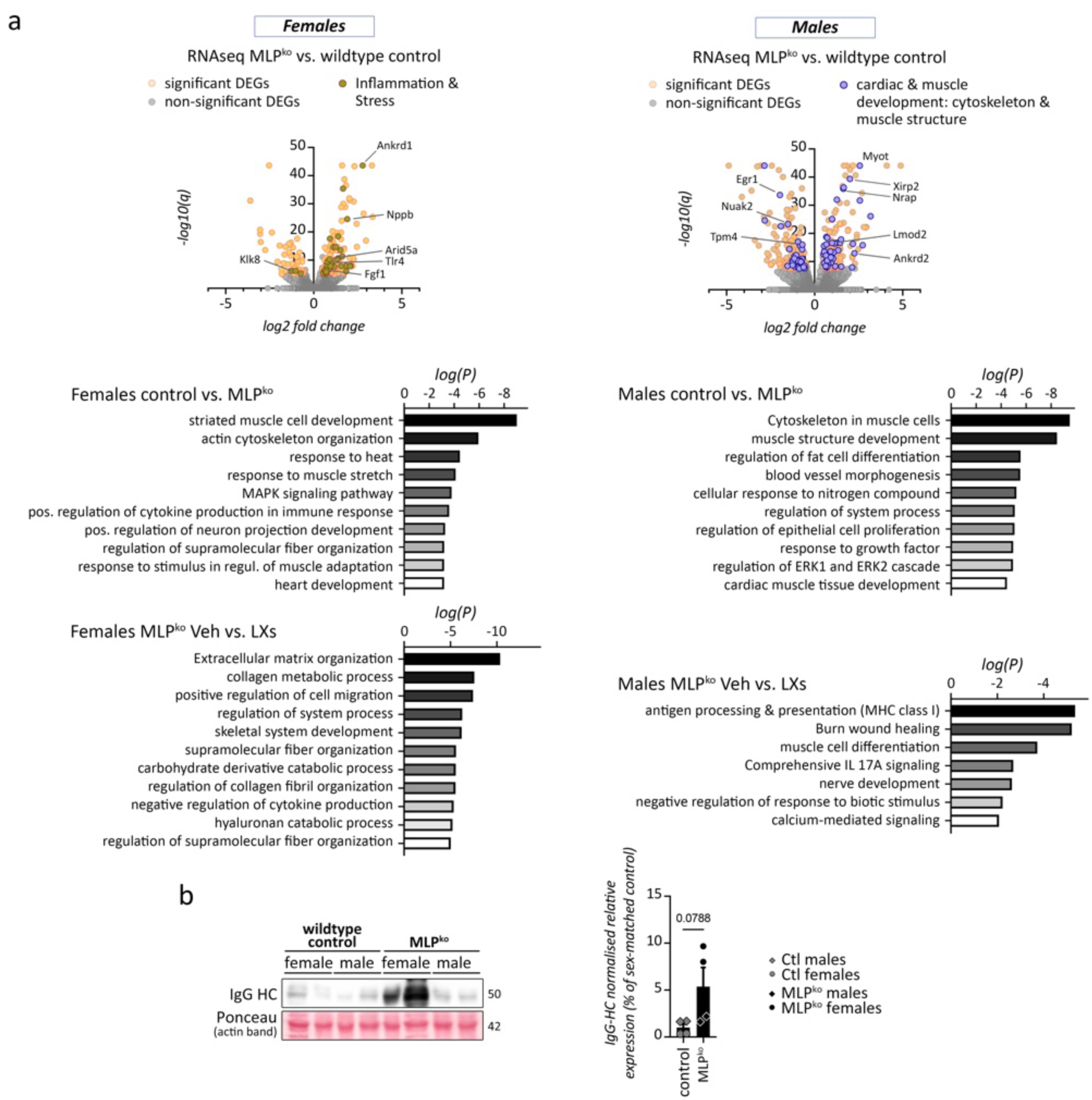
**a.** Enrichment analysis of significantly deregulated genes between male and female control and MLP^ko^ hearts (bottom left panels) and between male and female vehicle and lipoxin treated MLP^ko^ hearts (bottom right panels). Volcano plots (top) show select significantly deregulated genes belonging to the inflammation and stress (response to heat and positive regulation of cytokine production), as well as cardiac development (cytoskeleton and muscle structure) categories for wildtype vs. MLP^ko^ female and male hearts, respectively. **b.** Immunoblot analysis (left) and quantification (right) of immunoglobulin heavy chain (IgG HC) protein levels in cardiac extracts of male and female control and MLP^ko^ hearts. Data are presented as mean ± SEM (in % relative to sex-specific control). Sample sizes and p-values vs. control by unpaired Student’s t-test are indicated in the figure.

**Supplemental Figure S3.**
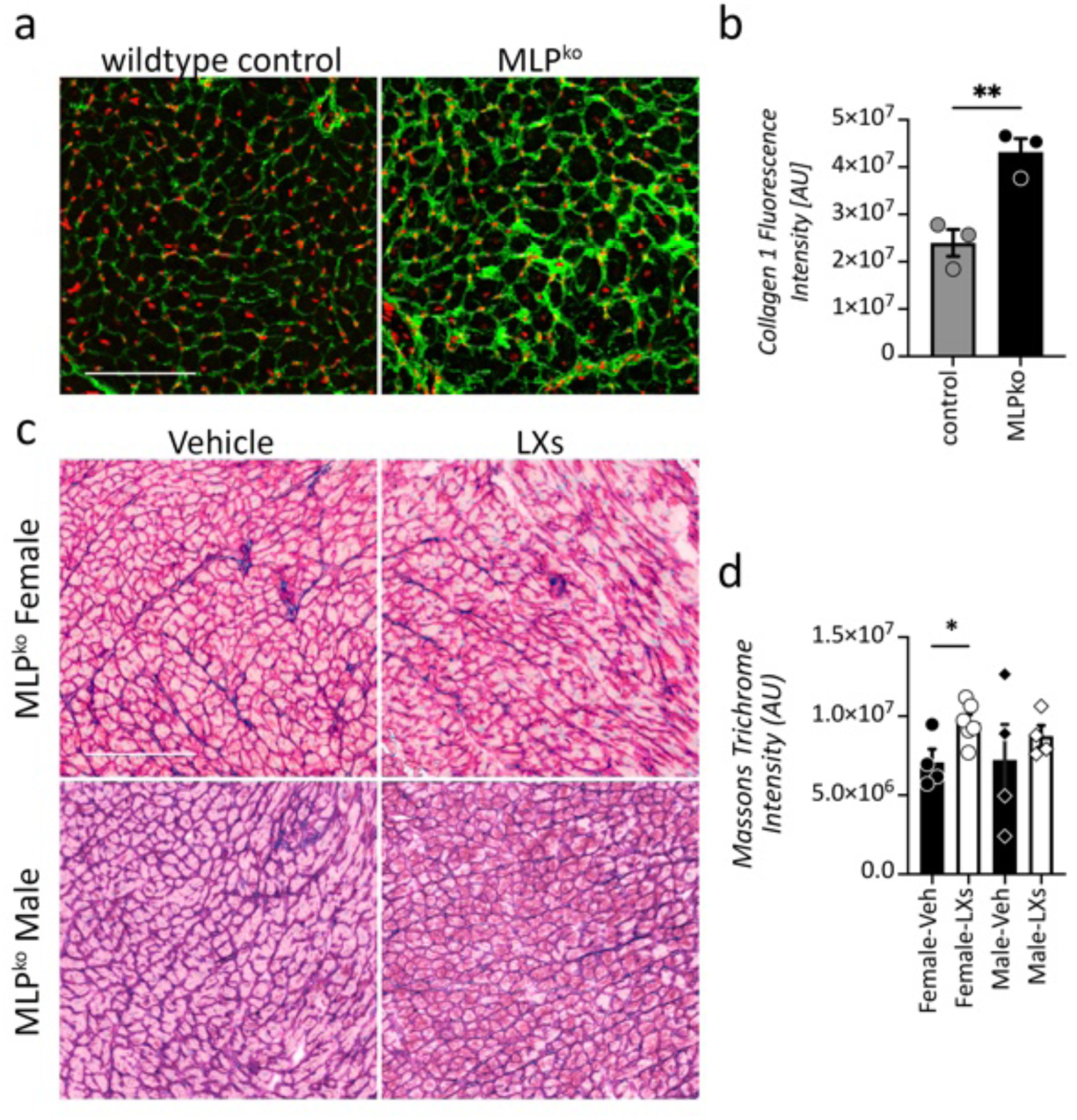
**a-b.** Immunofluorescence staining (a) and quantification (b) of collagen 1 staining (green) in frozen sections of control and MLP^ko^ hearts. DAPI (red) is used as counterstain. Data are presented as mean ± SEM. Sample sizes and p-values vs. control (** p<0.01) by unpaired Student’s t-test are indicated in the figure. Scalebar = 200µm. **c-d.** Masson trichrome stained cardiac sections of male and female MLP^ko^ mice treated with either vehicle or lipoxins (c) and quantification of staining intensity (d). Data are presented as mean ± SEM (in % relative to sex-specific vehicle treated MLP^ko^). Sample sizes and p-values vs. vehicle treated MLP^ko^ (* p<0.05) by Brown-Forsythe and Welch ANOVA using using Welch’s correction are indicated in the figure. Scalebar = 200µm.

**Supplemental Figure S4.**
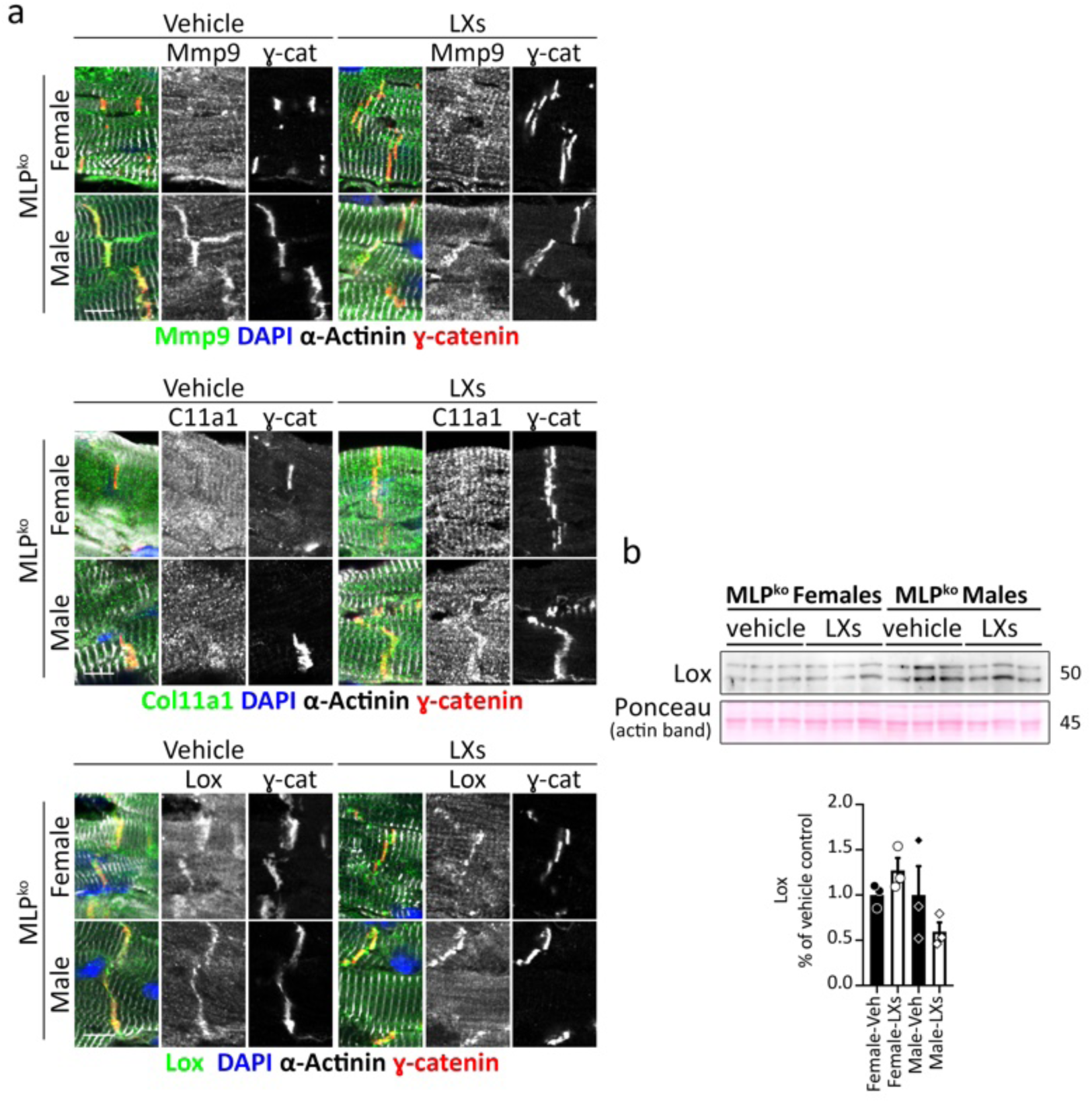
**a.** Immunofluorescence staining of Mmp9 (top panel, green in the overlay), Collagen 11a1 (C11a1, middle panel, green in the overlay) and Lox (bottom panel, green in the overlay) in frozen sections of vehicle or lipoxin treated male and female MLP^ko^ hearts. DAPI (blue), ɣ- catenin (ɣ-cat, red) and α-sarcomeric actinin-2 (white in the overlay) are used as counterstains. A set of different Mmp9, C11a1 and Lox antibodies was used compared to images shown in Figure 4a. Scalebar = 10µm. **b.** Immunoblot analysis (top) and quantification (bottom) of Lox protein levels in vehicle or lipoxin treated male and female MLP^ko^ hearts. Data are presented as mean ± SEM (in % relative to sex-specific vehicle treated MLP^ko^). Sample sizes and p-values vs. vehicle treated MLP^ko^ by Brown-Forsythe and Welch ANOVA using Welch’s correction are indicated in the figure.

**Supplemental Figure S5.**
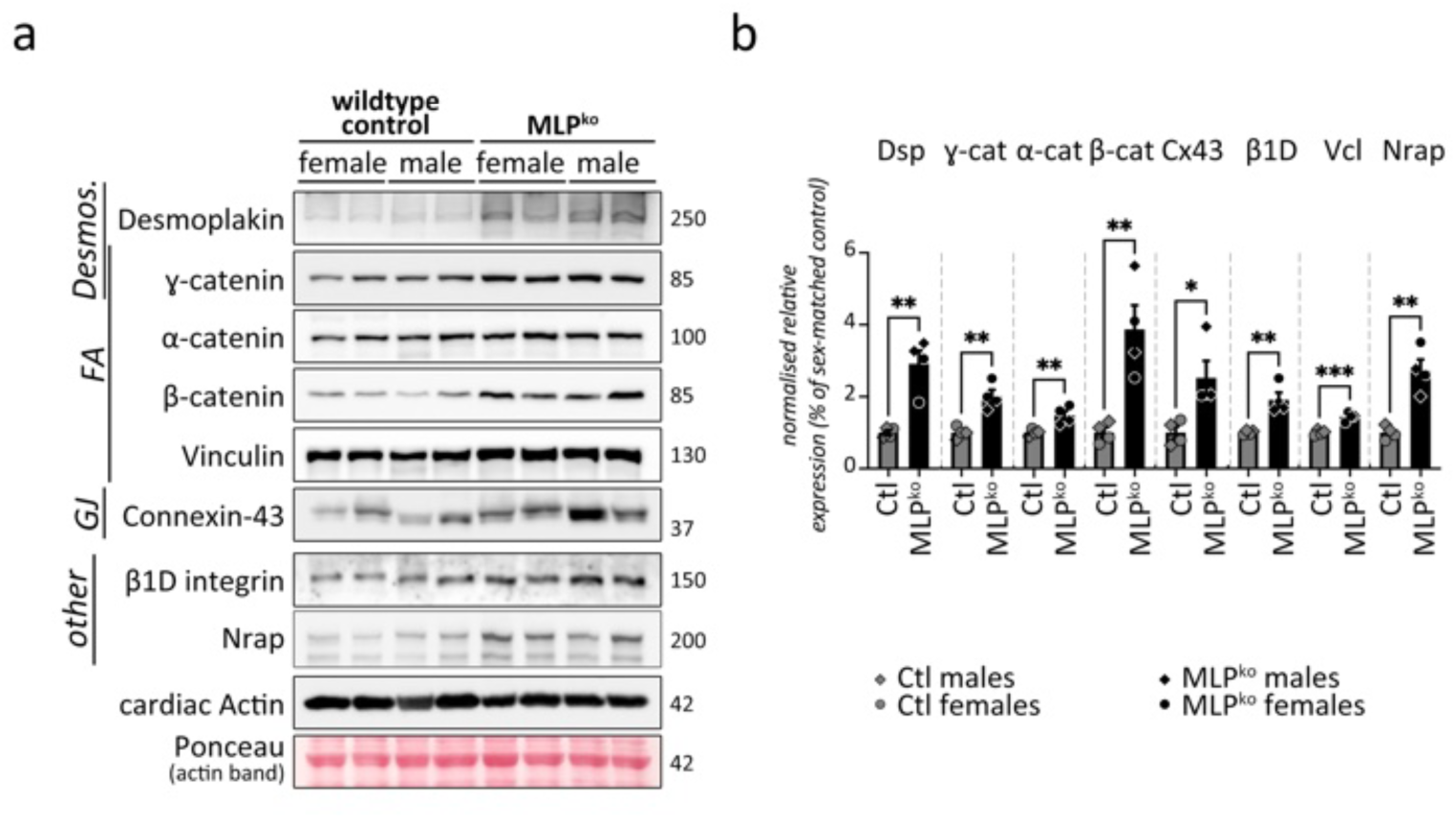
**a-b.** Immunoblot analysis (a) and quantification (b) of changes to intercalated protein expression levels in male and female control and MLP^ko^ hearts. Data are presented as mean values of normalized relative expression (of sex matched controls) ± SEM. Sample sizes and p-values vs. control (* p<0.05, ** p<0.01, *** p<0.001) by unpaired Student’s t-test are indicated in the figure. Abbreviations: Desmos. – desmosome, FA – facia adherens, GJ – gap junctions, DSP – desmoplakin, ɣ-cat - ɣ catenin, α-cat – α catenin, β-cat – β catenin, Cx43 – connexin-43, β1D – β1D integrin, Vcl – vinculin.

**Supplemental Figure S6.**
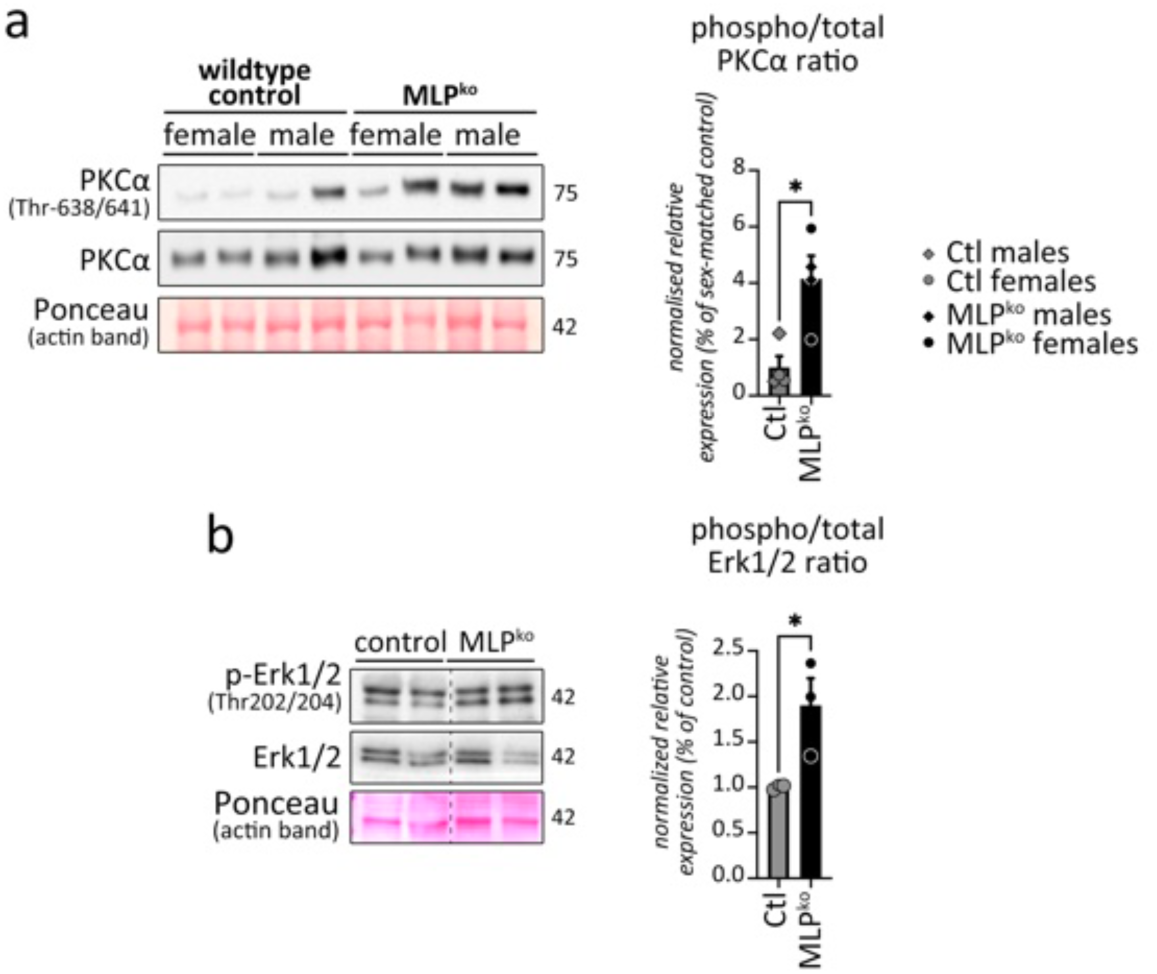
**a-b.** Immunoblot analysis (a) and quantification (b) of PKCα (a) and Erk1/2 (b) phosphorylation and total protein levels between male and female control and MLP^ko^ hearts. Ponceau is shown as loading control. Dashed lines in b indicate that bands were on the same blot, but in non- consecutive lanes. Data are presented as mean values of normalized relative expression (of sex matched controls in a) ± SEM. Sample sizes and p-values vs. control (* p<0.05) by unpaired Student’s t-test are indicated in the figure.

## References

1. Ahmed M, El Amrousy D, Hodeib H, Elnemr S (2023) Neutrophil-to-lymphocyte ratio as a predictive and prognostic marker in children with dilated cardiomyopathy. Cardiol Young 33:2493–2497 doi:10.1017/S1047951123000501

2. Arber S, Hunter JJ, Ross J, Jr., Hongo M, Sansig G, Borg J, Perriard JC, Chien KR, Caroni P (1997) MLP-deficient mice exhibit a disruption of cardiac cytoarchitectural organization, dilated cardiomyopathy, and heart failure. Cell 88:393–403 doi:10.1016/s0092-8674(00)81878-4

3. Avci A, Alizade E, Fidan S, Yesin M, Guler Y, Kargin R, Esen AM (2014) Neutrophil/lymphocyte ratio is related to the severity of idiopathic dilated cardiomyopathy. Scand Cardiovasc J 48:202–208 doi:10.3109/14017431.2014.932922

4. Baldeviano GC, Barin JG, Talor MV, Srinivasan S, Bedja D, Zheng D, Gabrielson K, Iwakura Y, Rose NR, Cihakova D (2010) Interleukin-17A is dispensable for myocarditis but essential for the progression to dilated cardiomyopathy. Circ Res 106:1646–1655 doi:10.1161/CIRCRESAHA.109.213157

5. Belkin AM, Zhidkova NI, Balzac F, Altruda F, Tomatis D, Maier A, Tarone G, Koteliansky VE, Burridge K (1996) Beta 1D integrin displaces the beta 1A isoform in striated muscles: localization at junctional structures and signaling potential in nonmuscle cells. J Cell Biol 132:211–226 doi:10.1083/jcb.132.1.211

6. Bermea KC, Duque C, Cohen CD, Bhalodia A, Rousseau S, Lovell J, Zita MD, Mugnier MR, Adamo L (2024) Myocardial B cells have specific gene expression and predicted interactions in dilated cardiomyopathy and arrhythmogenic right ventricular cardiomyopathy. Front Immunol 15:1327372 doi:10.3389/fimmu.2024.1327372

7. Borgeson E, Docherty NG, Murphy M, Rodgers K, Ryan A, O’Sullivan TP, Guiry PJ, Goldschmeding R, Higgins DF, Godson C (2011) Lipoxin A(4) and benzo-lipoxin A(4) attenuate experimental renal fibrosis. FASEB J 25:2967–2979 doi:10.1096/ñ.11-185017

8. Borgeson E, Johnson AM, Lee YS, Till A, Syed GH, Ali-Shah ST, Guiry PJ, Dalli J, Colas RA, Serhan CN, Sharma K, Godson C (2015) Lipoxin A4 Attenuates Obesity-Induced Adipose Inflammation and Associated Liver and Kidney Disease. Cell Metab 22:125– 137 doi:10.1016/j.cmet.2015.05.003

9. Borgeson E, McGillicuddy FC, Harford KA, Corrigan N, Higgins DF, Maderna P, Roche HM, Godson C (2012) Lipoxin A4 attenuates adipose inflammation. FASEB J 26:4287– 4294 doi:10.1096/ñ.12-208249

10. Canny GO, Lessey BA (2013) The role of lipoxin A4 in endometrial biology and endometriosis. Mucosal Immunol 6:439–450 doi:10.1038/mi.2013.9

11. Chandrasekharan JA, Sharma-Walia N (2015) Lipoxins: nature’s way to resolve inflammation. J Inflamm Res 8:181–192 doi:10.2147/JIR.S90380

12. Chen S, Wu RF, Su L, Zhou WD, Zhu MB, Chen QH (2014) Lipoxin A4 regulates expression of the estrogen receptor and inhibits 17beta-estradiol induced p38 mitogen-activated protein kinase phosphorylation in human endometriotic stromal cells. Fertil Steril 102:264–271 doi:10.1016/j.fertnstert.2014.03.029

13. Cohen L, Sagi I, Bigelman E, Solomonov I, Aloshin A, Ben-Shoshan J, Rozenbaum Z, Keren G, Entin-Meer M (2020) Cardiac remodeling secondary to chronic volume overload is attenuated by a novel MMP9/2 blocking antibody. PLoS One 15:e0231202 doi:10.1371/journal.pone.0231202

14. Dadson K, Kovacevic V, Rengasamy P, Kim GH, Boo S, Li RK, George I, Schulze PC, Hinz B, Sweeney G (2016) Cellular, structural and functional cardiac remodelling following pressure overload and unloading. Int J Cardiol 216:32–42 doi:10.1016/j.ijcard.2016.03.240

15. Ehler E, Horowits R, Zuppinger C, Price RL, Perriard E, Leu M, Caroni P, Sussman M, Eppenberger HM, Perriard JC (2001) Alterations at the intercalated disk associated with the absence of muscle LIM protein. J Cell Biol 153:763–772 doi:10.1083/jcb.153.4.763

16. Fu T, Mohan M, Brennan EP, Woodman OL, Godson C, Kantharidis P, Ritchie RH, Qin CX (2020) Therapeutic Potential of Lipoxin A4 in Chronic Inflammation: Focus on Cardiometabolic Disease. ACS Pharmacol Transl Sci 3:43–55 doi:10.1021/acsptsci.9b00097

17. Gibb AA, Lazaropoulos MP, Elrod JW (2020) Myofibroblasts and Fibrosis: Mitochondrial and Metabolic Control of Cellular Differentiation. Circ Res 127:427– 447 doi:10.1161/CIRCRESAHA.120.316958

18. Gilbert SJ, Wotton PR, Tarlton JF, Duance VC, Bailey AJ (1997) Increased expression of promatrix metalloproteinase-9 and neutrophil elastase in canine dilated cardiomyopathy. Cardiovasc Res 34:377–383 doi:10.1016/s0008-6363(97)00011-4

19. Giovannone N, Liang J, Antonopoulos A, Geddes Sweeney J, King SL, Pochebit SM, Bhattacharyya N, Lee GS, Dell A, Widlund HR, Haslam SM, Dimitroff CJ (2018) Galectin-9 suppresses B cell receptor signaling and is regulated by I-branching of N-glycans. Nat Commun 9:3287 doi:10.1038/s41467-018-05770-9

20. Guan J, Del Re DP (2025) Cell type specificity of Hippo-YAP signaling in cardiac development and disease. J Mol Cell Cardiol 207:51–63 doi:10.1016/j.yjmcc.2025.08.003

21. Halade GV, Lee DH (2022) Inflammation and resolution signaling in cardiac repair and heart failure. EBioMedicine 79:103992 doi:10.1016/j.ebiom.2022.103992

22. Harding D, Chong MHA, Lahoti N, Bigogno CM, Prema R, Mohiddin SA, Marelli-Berg F (2023) Dilated cardiomyopathy and chronic cardiac inflammation: Pathogenesis, diagnosis and therapy. J Intern Med 293:23–47 doi:10.1111/joim.13556

23. Hayward SL, Bautista-Lopez N, Suzuki K, Atrazhev A, Dickie P, Elliott JF (2006) CD4 T cells play major effector role and CD8 T cells initiating role in spontaneous autoimmune myocarditis of HLA-DQ8 transgenic IAb knockout nonobese diabetic mice. J Immunol 176:7715–7725 doi:10.4049/jimmunol.176.12.7715

24. Hinderer S, Schenke-Layland K (2019) Cardiac fibrosis - A short review of causes and therapeutic strategies. Adv Drug Deliv Rev 146:77–82 doi:10.1016/j.addr.2019.05.011

25. Hoque MM, Gbadegoye JO, Hassan FO, Raafat A, Lebeche D (2024) Cardiac fibrogenesis: an immuno-metabolic perspective. Front Physiol 15:1336551 doi:10.3389/fphys.2024.1336551

26. Huang Y, Huang LH, Su HB, Li YX, Chen H, Li JH, Yang LH, Su Q, Gui C (2025) Prognostic potential of neutrophil-to-lymphocyte ratio for adverse outcomes in dilated cardiomyopathy: a retrospective cohort study. Sci Rep 15:10339 doi:10.1038/s41598-025-94423-1

27. Ingle KA, Kain V, Goel M, Prabhu SD, Young ME, Halade GV (2015) Cardiomyocyte-specific Bmal1 deletion in mice triggers diastolic dysfunction, extracellular matrix response, and impaired resolution of inflammation. Am J Physiol Heart Circ Physiol 309:H1827–1836 doi:10.1152/ajpheart.00608.2015

28. Jaen RI, Fernandez-Velasco M, Terron V, Sanchez-Garcia S, Zaragoza C, Canales-Bueno N, Val-Blasco A, Vallejo-Cremades MT, Bosca L, Prieto P (2020) BML-111 treatment prevents cardiac apoptosis and oxidative stress in a mouse model of autoimmune myocarditis. FASEB J 34:10531–10546 doi:10.1096/ñ.202000611R

29. Jaen RI, Sanchez-Garcia S, Fernandez-Velasco M, Bosca L, Prieto P (2021) Resolution- Based Therapies: The Potential of Lipoxins to Treat Human Diseases. Front Immunol 12:658840 doi:10.3389/fimmu.2021.658840

30. Jaen RI, Sanchez-Garcia S, Fernandez-Velasco M, Cuadrado I, de Las Heras B, Bosca L, Prieto P (2025) New Biomarkers in the Diagnosis and Prognosis of Dilated Cardiomyopathy: Pro-Resolving Lipids and miRNAs. Cells 14 doi:10.3390/cells14231916

31. Kadhi A, Mohammed F, Nemer G (2021) The Genetic Pathways Underlying Immunotherapy in Dilated Cardiomyopathy. Front Cardiovasc Med 8:613295 doi:10.3389/fcvm.2021.613295

32. Karra L, Haworth O, Priluck R, Levy BD, Levi-Schaffer F (2015) Lipoxin B(4) promotes the resolution of allergic inflammation in the upper and lower airways of mice. Mucosal Immunol 8:852–862 doi:10.1038/mi.2014.116

33. Knoll R, Hoshijima M, Hoffman HM, Person V, Lorenzen-Schmidt I, Bang ML, Hayashi T, Shiga N, Yasukawa H, Schaper W, McKenna W, Yokoyama M, Schork NJ, Omens JH, McCulloch AD, Kimura A, Gregorio CC, Poller W, Schaper J, Schultheiss HP, Chien KR (2002) The cardiac mechanical stretch sensor machinery involves a Z disc complex that is defective in a subset of human dilated cardiomyopathy. Cell 111:943–955 doi:10.1016/s0092-8674(02)01226-6

34. Kong P, Christia P, Frangogiannis NG (2014) The pathogenesis of cardiac fibrosis. Cell Mol Life Sci 71:549–574 doi:10.1007/s00018-013-1349-6

35. Laezza F, Gerber BR, Lou JY, Kozel MA, Hartman H, Craig AM, Ornitz DM, Nerbonne JM (2007) The FGF14(F145S) mutation disrupts the interaction of FGF14 with voltage-gated Na+ channels and impairs neuronal excitability. J Neurosci 27:12033– 12044 doi:10.1523/JNEUROSCI.2282-07.2007

36. Laezza F, Lampert A, Kozel MA, Gerber BR, Rush AM, Nerbonne JM, Waxman SG, Dib-Hajj SD, Ornitz DM (2009) FGF14 N-terminal splice variants differentially modulate Nav1.2 and Nav1.6-encoded sodium channels. Mol Cell Neurosci 42:90–101 doi:10.1016/j.mcn.2009.05.007

37. Lange S, Gehmlich K, Lun AS, Blondelle J, Hooper C, Dalton ND, Alvarez EA, Zhang X, Bang ML, Abassi YA, Dos Remedios CG, Peterson KL, Chen J, Ehler E (2016) MLP and CARP are linked to chronic PKCalpha signalling in dilated cardiomyopathy. Nat Commun 7:12120 doi:10.1038/ncomms12120

38. Laroumanie F, Douin-Echinard V, Pozzo J, Lairez O, Tortosa F, Vinel C, Delage C, Calise D, Dutaur M, Parini A, Pizzinat N (2014) CD4+ T cells promote the transition from hypertrophy to heart failure during chronic pressure overload. Circulation 129:2111– 2124 doi:10.1161/CIRCULATIONAHA.113.007101

39. Lee C, Dartt DA (2024) Sex-dependent differential increase of specialized pro-resolving mediators in extracellular vesicles secreted by human primary conjunctival goblet cells during allergic inflammation. Life Sci 357:123058 doi:10.1016/j.lfs.2024.123058

40. Leoni G, Neumann PA, Sumagin R, Denning TL, Nusrat A (2015) Wound repair: role of immune-epithelial interactions. Mucosal Immunol 8:959–968 doi:10.1038/mi.2015.63

41. Li H, Zhu X, Cao X, Lu Y, Zhou J, Zhang X (2023) Single-cell analysis reveals lysyl oxidase (Lox)(+) fibroblast subset involved in cardiac fibrosis of diabetic mice. J Adv Res 54:223–237 doi:10.1016/j.jare.2023.01.018

42. Lou JY, Laezza F, Gerber BR, Xiao M, Yamada KA, Hartmann H, Craig AM, Nerbonne JM, Ornitz DM (2005) Fibroblast growth factor 14 is an intracellular modulator of voltage-gated sodium channels. J Physiol 569:179–193 doi:10.1113/jphysiol.2005.097220

43. Louis HA, Pino JD, Schmeichel KL, Pomies P, Beckerle MC (1997) Comparison of three members of the cysteine-rich protein family reveals functional conservation and divergent patterns of gene expression. J Biol Chem 272:27484–27491 doi:10.1074/jbc.272.43.27484

44. Lun AS, Chen J, Lange S (2014) Probing muscle ankyrin-repeat protein (MARP) structure and function. Anat Rec (Hoboken) 297:1615–1629 doi:10.1002/ar.22968

45. Lynch TLt, Ismahil MA, Jegga AG, Zilliox MJ, Troidl C, Prabhu SD, Sadayappan S (2017) Cardiac inflammation in genetic dilated cardiomyopathy caused by MYBPC3 mutation. J Mol Cell Cardiol 102:83–93

46. Ma ZG, Yuan YP, Wu HM, Zhang X, Tang QZ (2018) Cardiac fibrosis: new insights into the pathogenesis. Int J Biol Sci 14:1645–1657 doi:10.7150/ijbs.28103

47. Macdonald LJ, Boddy SC, Denison FC, Sales KJ, Jabbour HN (2011) A role for lipoxin A(4) as an anti-inflammatory mediator in the human endometrium. Reproduction 142:345–352 doi:10.1530/REP-11-0021

48. Maisch B, Alter P (2018) Treatment options in myocarditis and inflammatory cardiomyopathy : Focus on i. v. immunoglobulins. Herz 43:423–430 doi:10.1007/s00059-018-4719-x

49. Maisch B, Hufnagel G, Kolsch S, Funck R, Richter A, Rupp H, Herzum M, Pankuweit S (2004) Treatment of inflammatory dilated cardiomyopathy and (peri)myocarditis with immunosuppression and i.v. immunoglobulins. Herz 29:624–636 doi:10.1007/s00059-004-2628-7

50. Mauri M, Elli T, Caviglia G, Uboldi G, Azzi M (2017) RAWGraphs: A Visualisation Platform to Create Open Outputs. In: Proceedings of the 12th Biannual Conference on Italian SIGCHI Chapter. Association for Computing Machinery, Cagliari, Italy, p Article 28

51. Melinyte-Ankudavice K, Sukys M, Kasputyte G, Krikstolaitis R, Ereminiene E, Galnaitiene G, Mizariene V, Sakalyte G, Krilavicius T, Jurkevicius R (2024) Association of uncertain significance genetic variants with myocardial mechanics and morphometrics in patients with nonischemic dilated cardiomyopathy. BMC Cardiovasc Disord 24:224 doi:10.1186/s12872-024-03888-x

52. Myers MC, Berge A, Zhong Y, Maruyama S, Bueno C, Bastien A, Hofer K, Kaur R, Fazeli MS, Golchin N (2025) Prevalence and Incidence of Dilated Cardiomyopathy in the United States and Western Europe: A Systematic Review. Cardiol Res 16:295–305 doi:10.14740/cr2071

53. Nallanthighal S, Heiserman JP, Cheon DJ (2021) Collagen Type XI Alpha 1 (COL11A1): A Novel Biomarker and a Key Player in Cancer. Cancers (Basel) 13 doi:10.3390/cancers13050935

54. Ngwenyama N, Salvador AM, Velazquez F, Nevers T, Levy A, Aronovitz M, Luster AD, Huggins GS, Alcaide P (2019) CXCR3 regulates CD4+ T cell cardiotropism in pressure overload-induced cardiac dysfunction. JCI Insight 4 doi:10.1172/jci.insight.125527

55. Nigam S, Fiore S, Luscinskas FW, Serhan CN (1990) Lipoxin A4 and lipoxin B4 stimulate the release but not the oxygenation of arachidonic acid in human neutrophils: dissociation between lipid remodeling and adhesion. J Cell Physiol 143:512–523 doi:10.1002/jcp.1041430316

56. Pablo JL, Pitt GS (2017) FGF14 is a regulator of KCNQ2/3 channels. Proc Natl Acad Sci U S A 114:154–159 doi:10.1073/pnas.1610158114

57. Pablo JL, Wang C, Presby MM, Pitt GS (2016) Polarized localization of voltage-gated Na+ channels is regulated by concerted FGF13 and FGF14 action. Proc Natl Acad Sci U S A 113:E2665–2674 doi:10.1073/pnas.1521194113

58. Porter KE, Turner NA (2009) Cardiac fibroblasts: at the heart of myocardial remodeling. Pharmacol Ther 123:255–278 doi:10.1016/j.pharmthera.2009.05.002

59. Qi B, Wu QF, Yang ZJ, Huang N, Miao L (2025) Melatonin Attenuates Cardiac Dysfunction and Inflammation in Dilated Cardiomyopathy via M2 Macrophage Polarization. J Cardiovasc Pharmacol 85:156–165 doi:10.1097/FJC.0000000000001650

60. Russell R, Gori I, Pellegrini C, Kumar R, Achtari C, Canny GO (2011) Lipoxin A4 is a novel estrogen receptor modulator. FASEB J 25:4326–4337 doi:10.1096/ñ.11-187658

61. Schultheiss HP, Fairweather D, Caforio ALP, Escher F, Hershberger RE, Lipshultz SE, Liu PP, Matsumori A, Mazzanti A, McMurray J, Priori SG (2019) Dilated cardiomyopathy. Nat Rev Dis Primers 5:32 doi:10.1038/s41572-019-0084-1

62. Shai SY, Harpf AE, Babbitt CJ, Jordan MC, Fishbein MC, Chen J, Omura M, Leil TA, Becker KD, Jiang M, Smith DJ, Cherry SR, Loftus JC, Ross RS (2002) Cardiac myocyte-specific excision of the beta1 integrin gene results in myocardial fibrosis and cardiac failure. Circ Res 90:458–464 doi:10.1161/hh0402.105790

63. Sikking MA, Stroeks S, Henkens M, Venner M, Li X, Heymans SRB, Hazebroek MR, Verdonschot JAJ (2023) Cardiac Inflammation in Adult-Onset Genetic Dilated Cardiomyopathy. J Clin Med 12 doi:10.3390/jcm12123937

64. Sivakumar P, Gupta S, Sarkar S, Sen S (2008) Upregulation of lysyl oxidase and MMPs during cardiac remodeling in human dilated cardiomyopathy. Mol Cell Biochem 307:159–167 doi:10.1007/s11010-007-9595-2

65. Sun P, Zhang Z, Gao F, Yang C, Mang G, Fu S, Tian J, Chang J (2025) Silicate-based therapy for inflammatory dilated cardiomyopathy by inhibiting the vicious cycle of immune inflammation via FOXO signaling. Sci Adv 11:eadr7208 doi:10.1126/sciadv.adr7208

66. Turkowski K, Herzberg F, Gunther S, Brunn D, Weigert A, Meister M, Muley T, Kriegsmann M, Schneider MA, Winter H, Thomas M, Grimminger F, Seeger W, Savai Pullamsetti S, Savai R (2020) Fibroblast Growth Factor-14 Acts as Tumor Suppressor in Lung Adenocarcinomas. Cells 9 doi:10.3390/cells9081755

67. Vafiadaki E, Arvanitis DA, Sanoudou D (2015) Muscle LIM Protein: Master regulator of cardiac and skeletal muscle functions. Gene 566:1–7 doi:10.1016/j.gene.2015.04.077

68. van den Hoogen P, de Jager SCA, Huibers MMH, Schoneveld AH, Puspitasari YM, Valstar GB, Oerlemans M, de Weger RA, Doevendans PA, den Ruijter HM, Laman JD, Vink A, Sluijter JPG (2019) Increased circulating IgG levels, myocardial immune cells and IgG deposits support a role for an immune response in pre- and end-stage heart failure. J Cell Mol Med 23:7505–7516 doi:10.1111/jcmm.14619

69. van der Flier A, Gaspar AC, Thorsteinsdottir S, Baudoin C, Groeneveld E, Mummery CL, Sonnenberg A (1997) Spatial and temporal expression of the beta1D integrin during mouse development. Dev Dyn 210:472–486 doi:10.1002/(SICI)1097-0177(199712)210:4<472::AID-AJA10>3.0.CO;2-9

70. Wang J, Bai XQ, Li M, Wang XJ, Sun SW, Huang L, Zhang X, Chen X (2025) Comprehensive Plasma Oxylipin Profiling Reveals a Pro-Inflammatory Eicosanoid Signature and Diagnostic Biomarker Panel in Dilated Cardiomyopathy. Med Sci Monit 31:e950838 doi:10.12659/MSM.950838

71. Wang Q, McEwen DG, Ornitz DM (2000) Subcellular and developmental expression of alternatively spliced forms of fibroblast growth factor 14. Mech Dev 90:283–287 doi:10.1016/s0925-4773(99)00241-5

72. Wang S, Liu J, Wang M, Zhang J, Wang Z (2010) Treatment and prevention of experimental autoimmune myocarditis with CD28 superagonists. Cardiology 115:107–113 doi:10.1159/000256660

73. Wei EQ, Barnett AS, Pitt GS, Hennessey JA (2011) Fibroblast growth factor homologous factors in the heart: a potential locus for cardiac arrhythmias. Trends Cardiovasc Med 21:199–203 doi:10.1016/j.tcm.2012.05.010

74. Wu L, Ong S, Talor MV, Barin JG, Baldeviano GC, Kass DA, Bedja D, Zhang H, Sheikh A, Margolick JB, Iwakura Y, Rose NR, Cihakova D (2014) Cardiac fibroblasts mediate IL- 17A-driven inflammatory dilated cardiomyopathy. J Exp Med 211:1449–1464 doi:10.1084/jem.20132126

75. Xiong J, Zeng P, Cheng X, Miao S, Wu L, Zhou S, Wu P, Ye D (2013) Lipoxin A4 blocks embryo implantation by controlling estrogen receptor alpha activity. Reproduction 145:411–420 doi:10.1530/REP-12-0469

76. Yadav S, Sitbon YH, Kazmierczak K, Szczesna-Cordary D (2019) Hereditary heart disease: pathophysiology, clinical presentation, and animal models of HCM, RCM, and DCM associated with mutations in cardiac myosin light chains. Pflugers Arch 471:683–699 doi:10.1007/s00424-019-02257-4

77. Yan H, Pablo JL, Pitt GS (2013) FGF14 regulates presynaptic Ca2+ channels and synaptic transmission. Cell Rep 4:66–75 doi:10.1016/j.celrep.2013.06.012

78. Yang M, Fjaervoll HK, Fjaervoll KA, Wang NH, Utheim TP, Serhan CN, Dartt DA (2022) Sex-based differences in conjunctival goblet cell responses to pro-inflammatory and pro-resolving mediators. Sci Rep 12:16305 doi:10.1038/s41598-022-20177-9

79. Zhang J, Xu H, Li Z, Feng F, Wang S, Li Y (2025) Frequency of autoantibodies and their associated clinical characteristics and outcomes in patients with dilated cardiomyopathy: A systematic review and meta-analysis. Autoimmun Rev 24:103755 doi:10.1016/j.autrev.2025.103755

80. Zhou Y, Zhou B, Pache L, Chang M, Khodabakhshi AH, Tanaseichuk O, Benner C, Chanda SK (2019) Metascape provides a biologist-oriented resource for the analysis of systems-level datasets. Nat Commun 10:1523 doi:10.1038/s41467-019-09234-6

