## Supplemental File 1 (FACS gating strategy) for "Lipoxins Regulate Intercalated Disk-Associated Signaling and Immune Remodeling in Dilated Cardiomyopathy"

### MLP gating strategy + FMOs

### Gating strategy panel 1: Macrophages

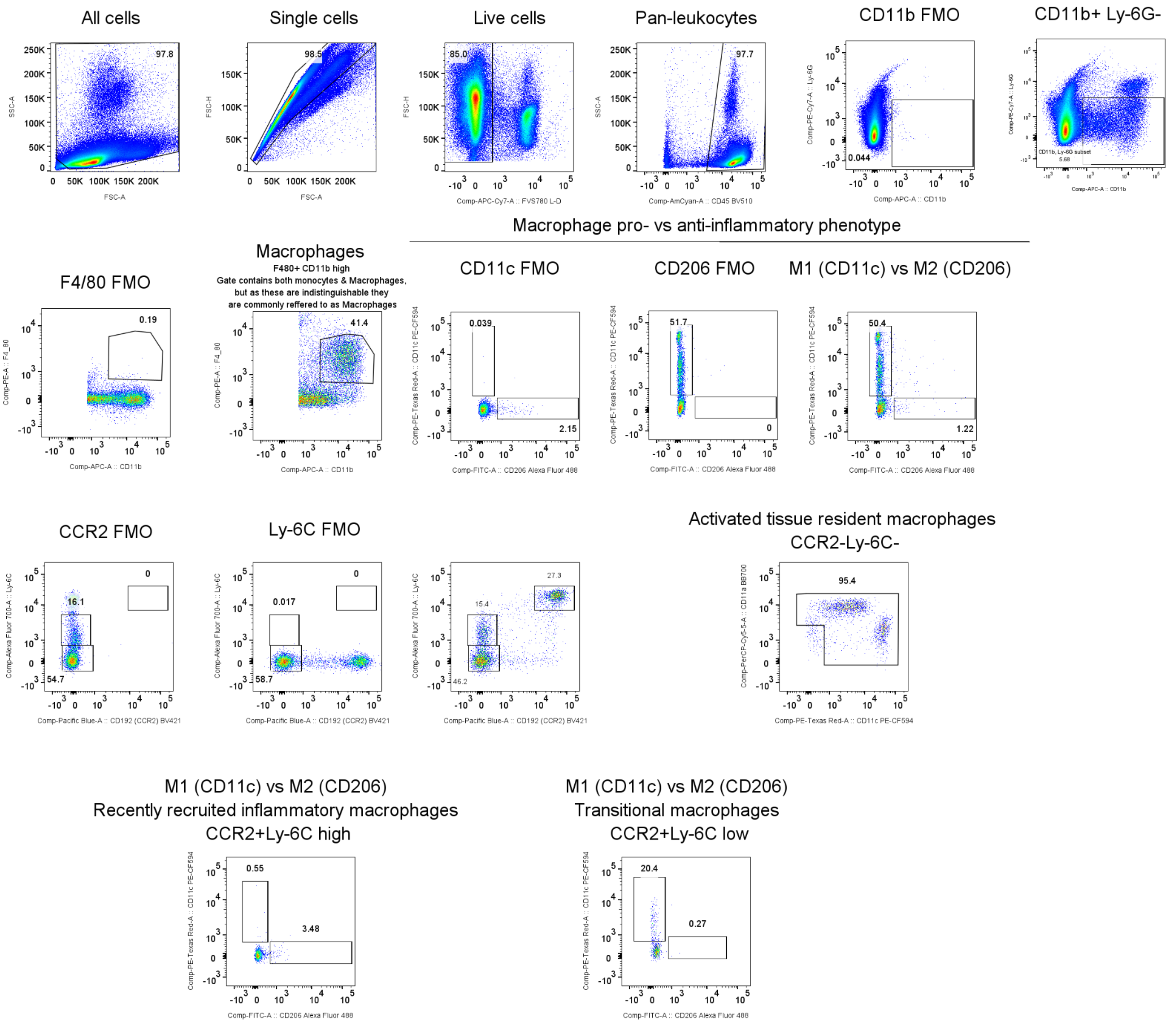

### Gating strategy panel 1: Neutrophils

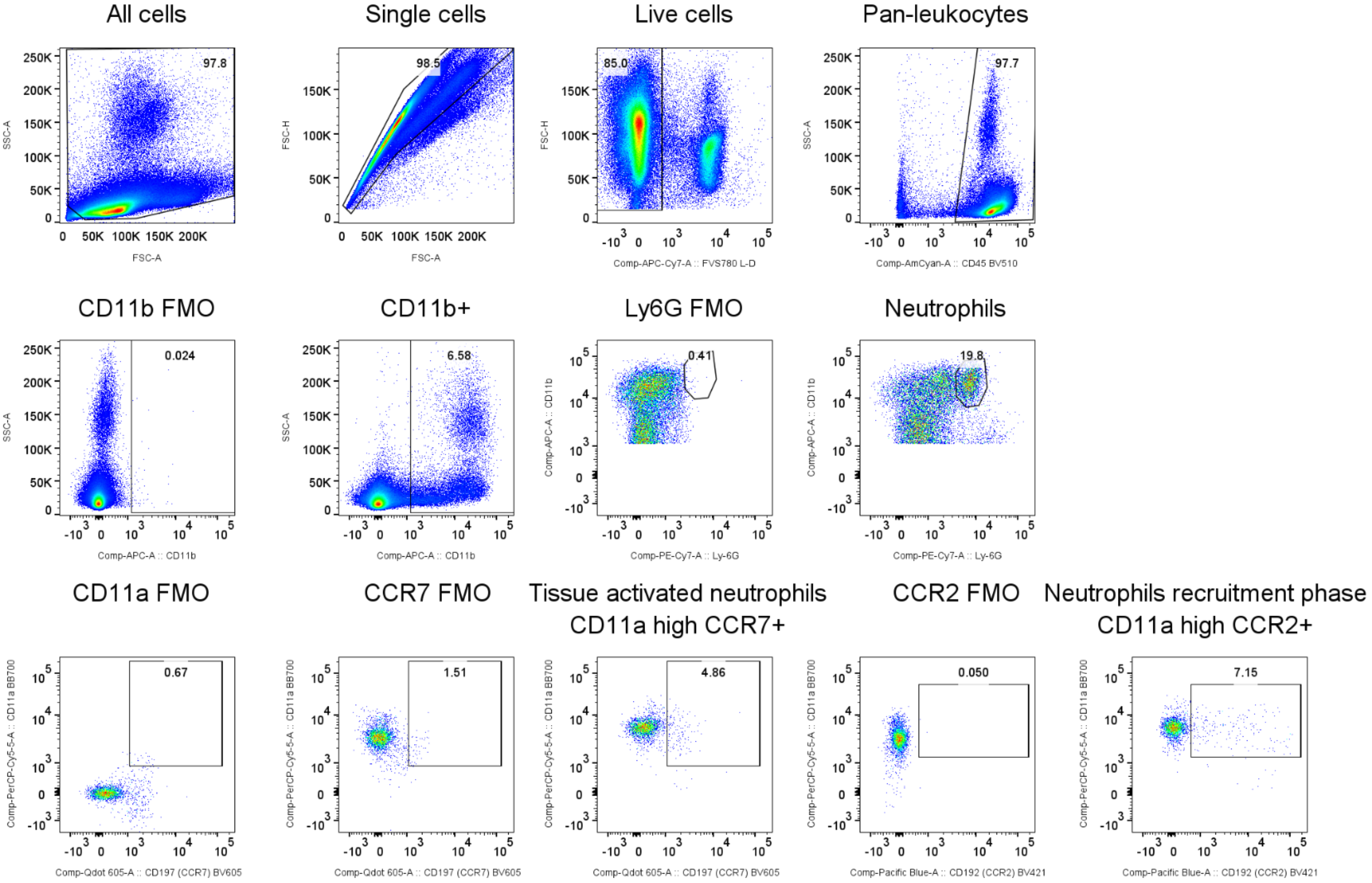

### Gating strategy panel 2: T- & B-cells

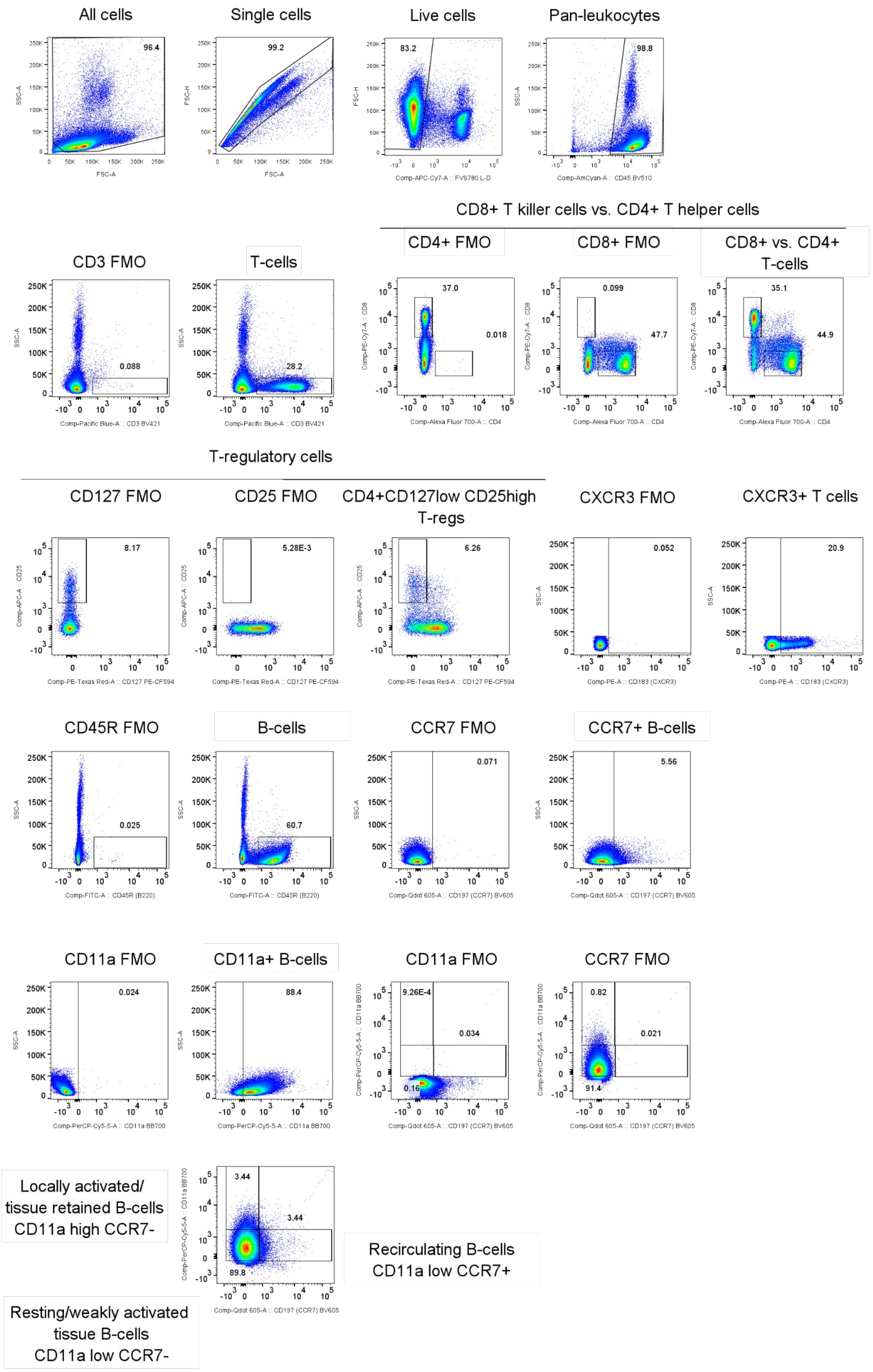
